# iClima: a ratiometric genetically encoded sensor for optical imaging of fast GABAergic chloride currents *in vitro* and *in vivo*

**DOI:** 10.64898/2026.05.27.728254

**Authors:** Giacomo Pasquini, Simone Giubbolini, Gabriele Nardi, Marco Brondi, Erko Beyene, Caterina Balsi, Alberto Egidi, Hanako Tsushima Semini, Silvia Landi, Marta Stancampiano, Andrew D Beale, Emily M Watson, Rachel S Edgar, John S O’Neill, Daniele Arosio, Melissa Santi, Anna Fassio, Claudia Lodovichi, Gian Michele Ratto

**Affiliations:** Institute of Biophysics, CNR, Pisa, Italy; National Enterprise for Nanoscience and Nanotechnology (NEST), Scuola Normale Superiore, Pisa, Italy; Department of Experimental Medicine, University of Genoa, Italy; Centre for Synaptic Neuroscience and Technology, Fondazione Istituto Italiano di Tecnologia, Genoa, Italy; Scuola Normale Superiore, Pisa, Italy; Istituto di Neuroscienze CNR, Pisa, Italy; Institute of Biophysics, CNR, Via alla Cascata 56/C, 38123 Trento, Italy; MRC Laboratory of Molecular Biology, Francis Crick Avenue, Cambridge CB2 0QH, UK; Department of Infectious Disease, Imperial College London, London W2 1NY, UK; Francis Crick Institute, 1 Midland Road, London, NW1 1AT, UK; IRCCS Azienda Ospedaliera Metropolitana, Genova, Italy

## Abstract

Fast GABAergic inhibition in the brain is mediated by chloride currents whose driving force depends on intracellular Chloride concentration ([Cl^−^]i). Resolving the spatiotemporal dynamics of [Cl^−^]i in neurons requires a sensor combining high Cl^−^ affinity, pH independence, and photostability - properties no existing tool achieves simultaneously. Here we introduce iClima (improved Chloride Imaging), a ratiometric genetically encoded sensor with high Cl^−^ affinity (K_d_=3.5 mM at pH 7.2), pH-insensitive under physiological conditions, and markedly improved photostability compared to existing sensors. These properties enable dynamic Cl^-^ imaging without simultaneous pH correction. We demonstrate that iClima resolves GABA_A_-driven Cl^−^ transients in cultured neurons, detects discrete Cl^−^ hotspots along dendrites, and captures sensory-evoked Cl^−^ transients *in vivo*. Finally, exploiting the spectral compatibility of iClima with GCaMP6f, we measure in interleaved trials Cl^−^ and Ca^2+^ transients in the same neurons *in vivo*, revealing a divergence between the somatic inhibitory drive and the neuronal excitation state as reported by Ca^2+^ dynamics.

## Introduction

Fast synaptic inhibition in the mature brain is primarily mediated by GABA and glycine, acting through ionotropic receptors permeable to chloride (Cl^−^). The driving force for these inhibitory currents depends on the intracellular Cl^−^ concentration ([Cl^−^]i): because the reversal potential of GABA_A_ and glycine receptors lies close to the resting membrane potential, even small fluctuations in [Cl^−^]i can shift inhibitory currents from hyperpolarising to depolarising, profoundly altering circuit function. The [Cl^−^]i landscape across a neuron is neither static nor uniform — different compartments receive specialised inhibitory inputs from distinct interneuron subtypes, and the rapid activation of inhibitory synapses, combined with the activity of Cl^−^ cotransporters KCC2 and NKCC1^1^, produces localised [Cl^−^]i transients that evolve continuously over time. This dynamic landscape, constitutes one element of ionic plasticity^2^, and shapes dendritic integration, network oscillations, and neuronal encoding. Yet, its spatiotemporal architecture remains poorly understood as measuring neuronal [Cl^−^]i with adequate resolution has proven technically challenging.

Genetically encoded fluorescent sensors have emerged as the most promising tools for imaging [Cl^−^]i in neurons. Among the most widely used are LSSmClopHensor^3–5^ and SuperClomeleon^6–11^, both of which have provided valuable insights into chloride homeostasis. However, both suffer from critical limitations. LSSmClopHensor has low Cl^−^ affinity at physiological pH (K_d_∼57 mM at pH 7.2), poorly matched to the physiological [Cl^−^]i of mature neurons (5–10 mM). SuperClomeleon improves affinity (20–40 mM) but its FRET-based readout entangles Cl^−^ and pH signals, precluding quantitative measurement without simultaneous pH correction. Crucially, both sensors exhibit strong pH sensitivity within the physiological range, meaning that pH fluctuations accompanying neuronal activity can introduce substantial artefacts^12^. Together, these shortcomings — low affinity, pH sensitivity, and insufficient photostability — have prevented reliable measurement of Cl^−^ transients under physiological conditions *in vivo*.

Here we introduce iClima (improved Chloride Imaging), a ratiometric genetically encoded sensor designed to overcome these limitations, and demonstrate its ability to resolve Cl^−^ transients *in vitro* and *in vivo*.

## Results

### iClima design

The new sensor iClima (improved Cl^−^ imaging) is a fusion between mCRISPRed, a pH- and Cl^−^-insensitive red fluorescent protein, and mClover3*, an engineered green mutant of mClover3. As in the closely related sensor LSSmClopHensor^3,4^, the green moiety serves as the Cl^−^-sensing element while the red moiety provides a ratiometric reference that is analyte-insensitive. Protonation of residues near the green fluorophore is required for anion access to the binding pocket, making Cl^−^ measurements intrinsically pH-dependent. Consequently, most genetically encoded Cl^−^ sensors require simultaneous pH measurement to yield reliable estimate of [Cl^−^]i.

The development of mClover3* was guided by two objectives: (i) to shift the proton affinity sufficiently high that the sensor remains nearly fully protonated across the physiological pH range, thereby decoupling Cl^−^ measurements from pH fluctuations; and (ii) enhance Cl^−^ affinity to maximise sensitivity under physiological conditions. We selected pK_a_ values above the physiological range,to ensure near-complete protonation, and therefore maximal Cl^−^ sensitivity, at resting intracellular pH, while preserving or enhancing fluorescence brightness.

mClover3 was chosen as the mutagenesis scaffold owing to its exceptional brightness and photostability, properties previously shown to confer advantages for chloride sensing applications^13,14^. The resulting mutant, mClover3*, is linked to mCRISPRed via a flexible amino acid linker (Figure 1a). mCRISPRed^15^ was selected from a panel of long-Stokes-shift red fluorescent proteins evaluated for pH insensitivity (pKa<5.5), photostability, and sufficent spectral separation from mClover3* to minimise FRET.

**Figure 1.**
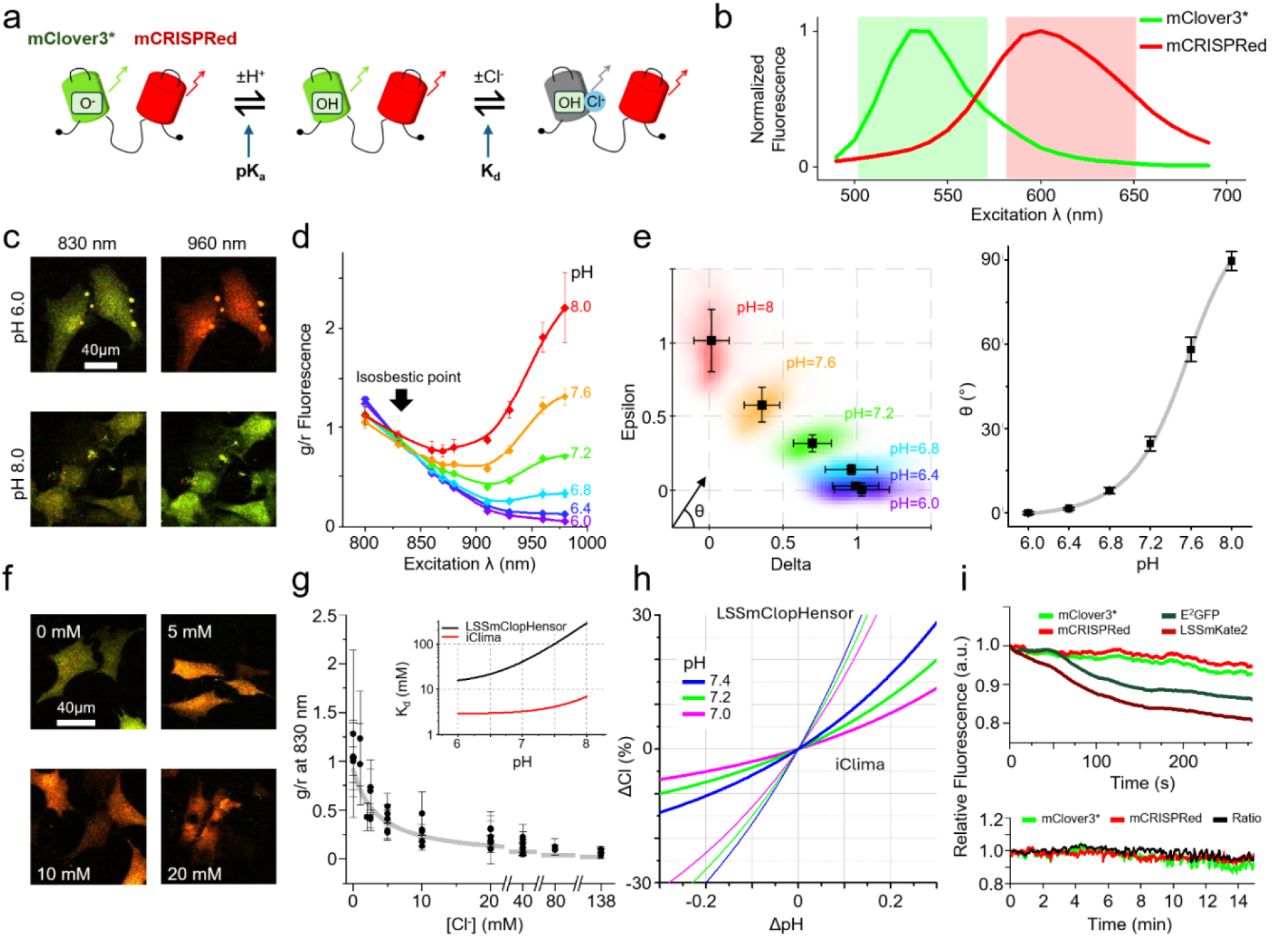
Structure and spectral properties of iClima *in vitro*. **a**) iClima is mostly protonated at physiological pH and fluorescence is quenched by Cl^−^. **b**) mClover3* and mCRISPRed emission spectra excited at 450 nm at pH 7.2 (262 and 61 cells for mCRISPRed and mClover3*, respectively). The red and green areas indicate the emission filter bandpass used in dual-channel imaging. **c**) Two-photon imaging of iClima in MEF cells at pH 6 and pH 8 in presence of ionophores, excited at two excitation wavelengths. **d**) Dependence of two-photon excitation spectra on pH. Green fluorescence (g) is normalised to red fluorescence (r). Data corrected for bleed-through (3 plates for each pH, 798 cells in total). **e**) Decomposition of the excitation spectra as a linear combination of the spectra obtained at pH 6 and pH 8. The angle θ formed by a vector joining each cell with the origin provides an estimate of pH. The right panel shows the calibration curve provided by Eq. 1, that delivers the *pKa*; (sample size as in d). **f**) Two-photon imaging of MEF cells at different [Cl^−^]i. **g**) The average g/r ratio from 6 calibration rounds was fitted to obtain iClima’s *Kd* (6 calibration rounds with an average of 4 points for each [Cl^-^], 40 measurements in total). Inset: comparison of the apparent *Kd’* for iClima and LSSmClopHensor. **h**) Dependence of the error in [Cl^−^]i measurement on the error in pH measurement, at different pH values. **i**) Top: photobleaching rates of iClima components compared to LSSmClopHensor (2 and 1 mouse and 24 and 10 cells respectively for iClima and LssmClopHensor). Bottom: baseline two-photon iClima acquisition in mouse pyramidal cells in vivo (2 frames per second, 130 µm depth; 65 cells from 1 mouse). Fluorescence was corrected for dark offset and bleed-through.

### Spectroscopic characterisation of iClima

The emission spectra of the two proteins are shown in Fig. 1b. Depending on the specifics of the emission filters, there is some bleed-through of mCRISPRed into the green channel and, to a lesser extent, of mClover3* into the red channel. These values are specific to the imaging set-up and must be determined beforehand; all data must be corrected using these factors (see Methods and Supp. Fig. 2).

**Figure 2.**
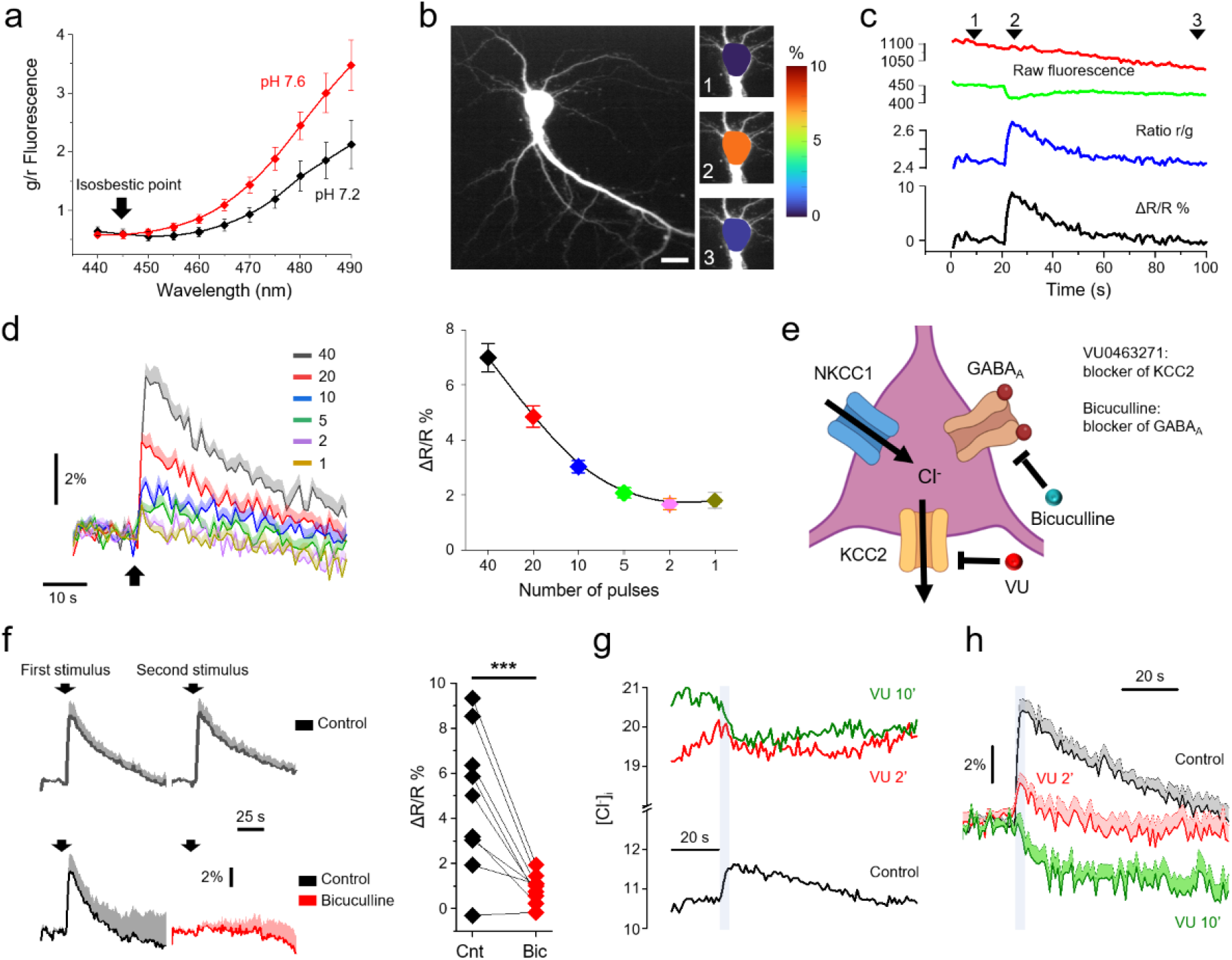
Cl^−^ transients in primary mouse hippocampal neurons. **a**) One-photon excitation spectra of iClima. **b)** Representative image of a neuron. Insets show the Cl^−^ change measured at three time points indicated by arrowheads in panel c. **c)** Raw fluorescence (red and green channels), r/g ratio (blue), and ΔR/R percentage values in response to extracellular stimulation (40 depolarising pulses, 1 ms each, 20 Hz). **d)** Responses to stimuli comprising an increasing number of depolarising pulses (n=18 neurons). Black arrows indicate stimulus onset. **e)** Schematic illustration of the main drivers of intracellular Cl^−^ dynamics: the cotransporters NKCC1 (Cl^−^ influx) and KCC2 (Cl^−^ efflux), and the ionotropic GABAA receptor. Bicuculline is a competitive GABAA antagonist; VU (VU0463271) is a KCC2 inhibitor. Generated with BioRender.com. **f**) Responses to two successive stimuli (40 pulses each, 5 min apart) under control conditions (top, n=13 cells) or in the presence of 30 µM bicuculline (bottom, n=9 cells). Solid line: mean; shaded area: SEM. Response amplitudes did not differ significantly between the two successive stimuli under control conditions (paired Wilcoxon signed-rank test, two-sided, p=0.88). Bicuculline suppressed the response (right panel; paired Wilcoxon signed-rank test two sided, p<0.006). **g**) Baseline [Cl^−^]i of a representative neuron recorded under control conditions (black), after 2 min (red), and after 10 min (green) of VU0463271 treatment (0.5 µM). Baseline [Cl^−^]i increased progressively; shaded bar indicates the stimulation period. A clear increase of baseline [Cl^−^]i during VU0463271 treatment was observed in 5/12 neurons. **h**) Kinetics of Cl^−^ transients upon stimulation (shaded area) across all analysed cells under control conditions and after VU0463271 treatment. Solid line: mean; shaded area: SEM. ΔR/R normalised to baseline before the stimulus (Control and 2 min VU, n=12; 10 min VU, n=9).

We transfected murine embrionic fibroblasts (MEF) with iClima and used ionophores permeable to H^+^ and Cl^−^ to control intracellular pH and [Cl^−^]i. Cells were imaged in zero Cl^−^ at pH values ranging from 6.0 to 8.0 using two-photon excitation at several excitation wavelengths (Fig. 1c,d). The signals collected in the green and red channels were dark-subtracted and corrected for bleed-through. In the following, g and r indicate the fluorescece of mClover3* and mCRISPRed respectively. To compensate for fluctuations in laser intensity and for power changes at different wavelengths, all spectra were computed as the g/r ratio between the signals from mClover3* and mCRISPRed (Fig. 1d). The excitation spectra of iClima are dependent on pH: the deprotonated sensor peaks at wavelengths >980 nm, while at 830 nm the fluorescence is independent of pH (Fig. 1c,d). The excitation spectrum (g/r ratio) X(λ) of a cell can be expressed as a linear combination of the average excitation spectra at pH 6.0 and 8.0 (X_pH=6_ and X_pH=8_), with X(λ) = δX_pH=6_(λ) + εX_pH=8_(λ). The angle described by δ and ε (Fig. 1e) depends on pH according to Eq. 1. By fitting the calibration data with Eq. 1 we obtained a *pKa* of 7.84±0.01 (Fig. 1e).

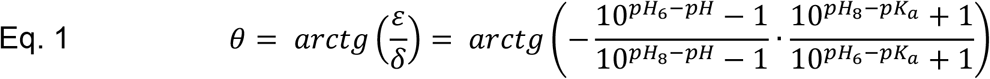

Cl^−^ calibration was performed at pH 7.2 by varying the Cl^−^ concentration of the medium in the presence of ionophores. As expected, the fluorescence of mClover3* decreased with increasing [Cl^−^] (Fig. 1f), and Figure 1g shows the dependence of the g/r ratio on [Cl^−^]. The calibration curve was obtained from 6 runs performed in Pisa, Cambridge and Genoa using both 1-photon and 2-photon excitation, a multi-laboratory validation that substantially strengthens confidence in the calibration. On average, about 600 cells were imaged per each Cl^−^ concentration. Data from each dataset were first divided by the median value obtained at [Cl^−^] = 0 mM to normalize for setup-specific factors, then pooled and fitted to Eq. 2, which reads:

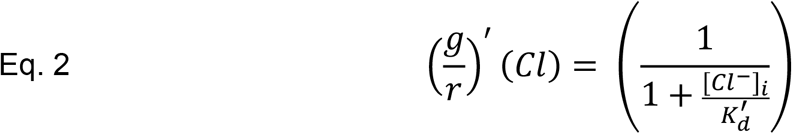

where 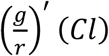 is the green/red ratio normalized by the mean 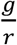 at Cl=0 mM and 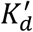 is the affinity corrected for the fraction of Cl sensitive protein at a given pH.

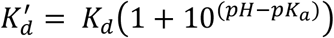

The fit (Fig. 1g) returned an apparent dissociation constant for Cl^−^ of 3.5±0.8 mM at pH 7.2, indicating that iClima’s affinity for Cl^−^ is roughly 15-fold higher than that of its predecessor LSSmClopHensor, whose affinity at pH 7.2 was 56.9 mM. The inset of Fig. 1g shows that the apparent affinity of iClima for Cl^−^ is far less dependent on pH than that of LSSmClopHensor.

Accordingly, estimates of [Cl^−^]i obtained with iClima are substantially less affected by pH errors: an undetected pH change of 0.1 units causes an error in the [Cl^−^]i estimate of approximately 4% with iClima, compared to ∼20% with LSSmClopHensor (Fig. 1h). Therefore, dynamic Cl^-^ measurement under physiological conditions could be captured by iClima imaging performed at a single excitation wavelength, as neuronal pH doesn’t substantially deviate from basal levels (7.1 - 7.2).

Since dynamic imaging requires high resistance to photobleaching, we measured the photostability of mClover3* and mCRISPRed under prolonged two-photon excitation at the isosbestic wavelength (830 nm) and compared it to LSSmClopHensor. The photobleaching time constants of iClima components were substantially longer than those of LSSmClopHensor (τ=72 min for mClover3* vs. 28 min for E^2^GFP; τ=105 min for mCRISPRed vs. 22 min for LSSmKate2; Fig. 1i, top). Continuous two-photon imaging of pyramidal neurons expressing iClima over 15 min confirms that prolonged *in vivo* acquisition is feasible with minimal photobleaching (Fig. 1i, bottom).

### Dynamic Cl^-^ imaging in cultured neurons

To assess whether iClima can report the dynamics of GABA_A_-mediated chloride fluxes, we transfected primary hippocampal neurons with the sensor at 14 DIV and imaged at the 1-photon isosbestic wavelength of iClima from 17 DIV (Fig. 2a). Neurons were stimulated with 1 ms depolarising current pulses at 20 Hz delivered through extracellular platinum–iridium electrodes. Glutamatergic transmission was suppressed with D-APV and CNQX, and cells were maintained at 30°C throughout imaging. Acquisition was performed on a wide-field microscope by alternating the two emission channels using a high-speed filter wheel (Fig. 2b). Fluorescence from the two channels was corrected for offset and bleed-through; the ratio R of the fluorescence emitted by the two moieties of iClima provided an estimate of the Cl^−^ change relative to baseline (Eq. 3, Fig. 2c), with CRISPRed fluorescence as the numerator so that R increases with increasing [Cl^−^]i.

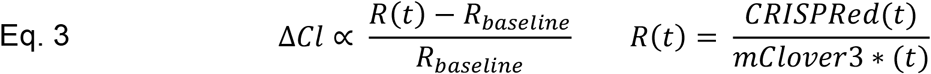

Field stimulation evoked clear responses whose amplitude correlated with the number of depolarising pulses (Fig. 2d). In principle, the observed changes (approximately 6% for the largest transient) could reflect cellular acidification rather than a Cl^−^ increase. However, based on the pH calibration (Fig. 1d), such a response would require an acidification of nearly 0.2 pH units — a magnitude attained only during prolonged seizures^4^ and therefore unrealistic under our stimulation conditions. To confirm the GABAergic origin of the responses, we blocked GABA_A_ receptors with bicuculline (Fig. 2f). Having verified that responses to two sequential stimuli were reproducible, bicuculline was sufficient to suppress the second response, confirming the obligatory role of GABA_A_ receptor activation. The recovery of [Cl^−^]i to baseline following stimulation follows an exponential time course with a median time constant of 42 s (Median Absolute Deviation, MAD=13.8), substantially slower than recovery time constants measured *in vivo* (Fig. 4f; Supp. Fig. 3), consistent with the known delayed functional maturation of KCC2 in dissociated hippocampal cultures^16^. Finally, we tested the dependence of the response on the Cl^−^ gradient by inhibiting KCC2 with VU0463271 (0.5 µM). Within 2 minutes of treatment baseline [Cl^−^]i rose progressively (Fig. 2g); after 10 minutes, the response to extracellular stimulation reversed polarity, indicating that GABA had become depolarising, consistent with Cl^−^ accumulation following KCC2 blockade (Fig. 2g,h). Together, these pharmacological manipulations confirm that iClima faithfully reports [Cl^−^]i dynamics driven by GABA_A_ receptor activation.

**Figure 3.**
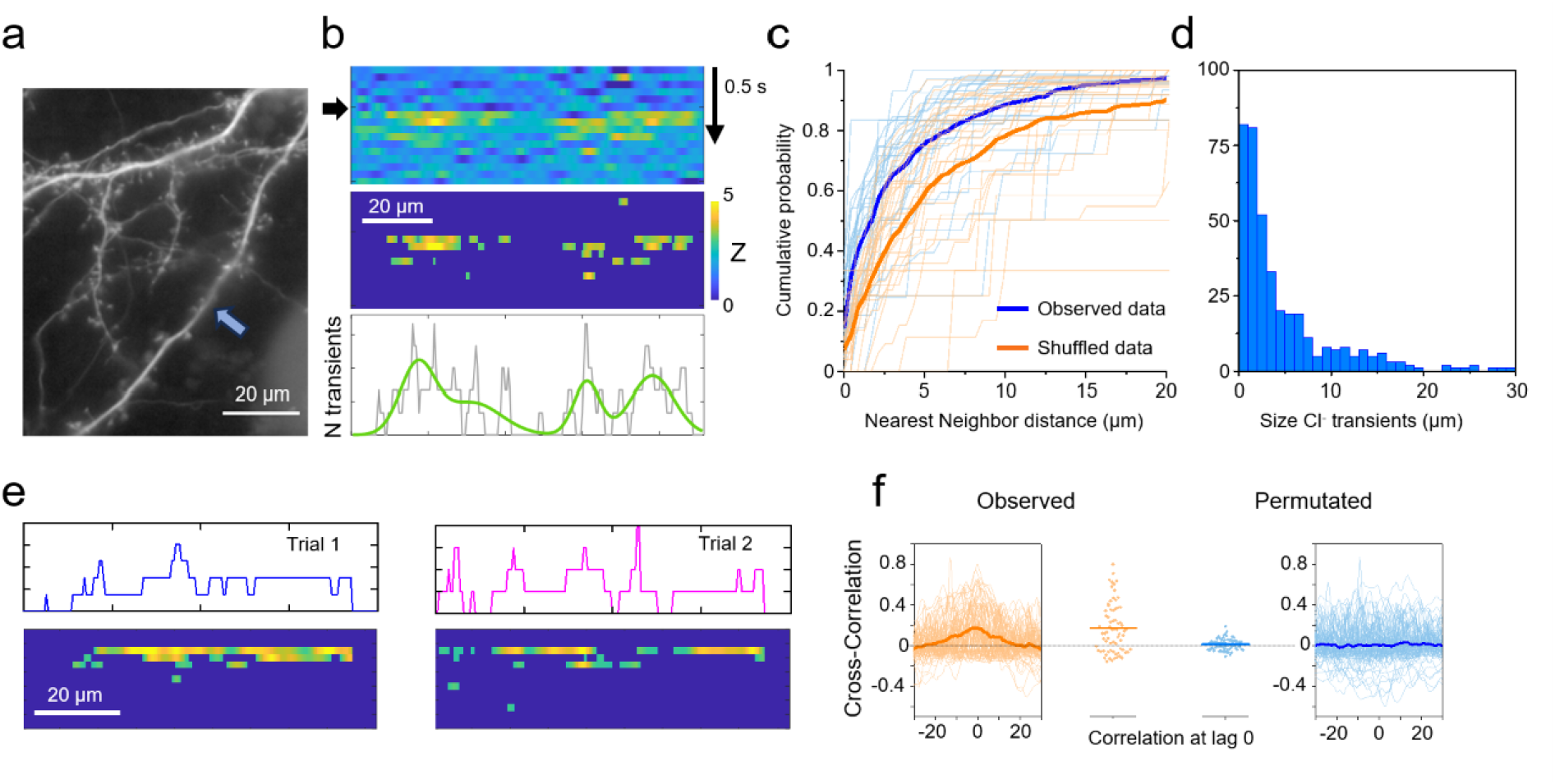
Subcellular spatial distribution of Cl^−^ hotspots. **a**) Wide-field image in which the arrow indicates the dendrite analysed in the right panels. **b**) Top: derivative of the fluorescence measured in the green channel as expressed by Eq. 4. Middle: Z-score of the derivative thresholded at Z<3. Bottom: density of Cl^−^ transients as a function of position along the dendrite; the green curve shows the Gaussian mixture model fit used to estimate hotspot size. **c**) The blue line shows the cumulative distribution of nearest-neighbour distances across all regions where the derivative of the Cl^−^ transient exceeded threshold; thin lines indicate the distribution for each individual dendrite (28 dendrites, 3 acquisition rounds). The orange line indicates the nearest-neighbour distribution obtained by placing all transients at random positions along each dendrite while preserving their size and number. The observed distribution was markedly shifted to the left relative to the randomised data (two sided Kolmogorov-Smirnov test, p<<0.001), indicating non-random clustering. **d**) Distribution of identified hotspot sizes expressed as full width at half maximum of the Gaussian model. **e**) Cl^-^ hotspots observed in two different trials on the same dendrite and their density profiles. **f**) Cross-correlation of the hotspot density profiles across repeated trials (n=25 dendrites). The central peak indicates that Cl^−^ hotspots recur at consistent dendritic locations across trials (left, orange). Symbols indicate the cross-correlation at lag 0 for all pairs of repeated trials. After random circular permutation of the density profiles the central peak disappears, confirming that spatial overlap is not due to chance.

**Figure 4.**
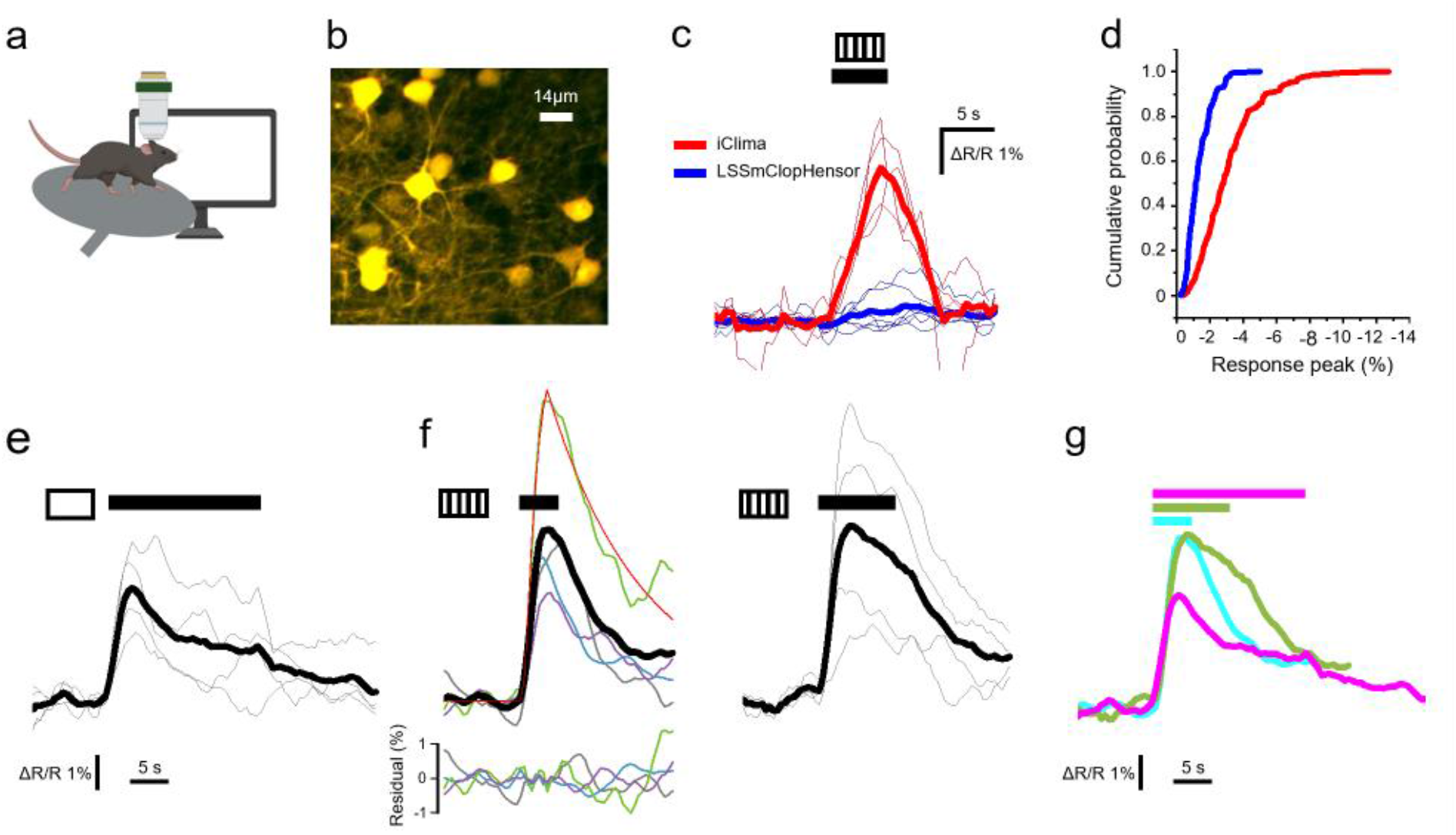
Chloride responses to visual stimuli. **a**) Schematic of the experiment. Mice were habituated to head fixation under the two-photon microscope and imaged in the awake head-restrained condition. Illustration generated with BioRender.com. **b**) Representative two-photon field of view showing iClima-expressing pyramidal neurons in V1 **c**) Comparison of Cl^−^ responses recorded in iClima- and LSSmClopHensor-expressing pyramidal neurons under identical visual stimulation (5 s drifting grating). Thin lines: mean response per animal (n=4 iClima mice, 366 neurons; n=7 LSSmClopHensor mice, 714 neurons); thick lines: grand mean across animals. The near-absence of a detectable response with LSSmClopHensor confirms the superior Cl^−^ sensitivity of iClima and rules out pH and haemodynamic artefacts. **d**) Cumulative distribution of response amplitudes from 189 (LSSmClopHensor) and 245 (iClima) imaging trials. **e**) Mean Cl^−^ responses of pyramidal neurons in isoflurane- anaesthetised mice to a full-field luminance step lasting 20 s (n=4 mice, 20 sessions, 817 neurons). Thin lines: mean response per mouse; thick line: grand mean. Horizontal bar indicates stimulus timing. **f**) Cl^−^ transients in response to a drifting grating lasting 5 s (left) or 10 s (right). Thick line: mean across mice (n=4); thin lines: mean responses for individual mice (5 s grating: 10 sessions, 607 neurons; 10 s grating: 12 sessions, 238 neurons). The red line shows the piecewise logistic plus exponential fit; the bottom panel shows the fit residuals across all four mice (two-sided t-test, mean=0, p>0.80). Horizontal bars indicate stimulus timing. **g)** Overlay of mean Cl^−^ responses to the three stimuli shown in panels e and f. Magenta: full-field luminance step; green: 10 s drifting grating; cyan: 5 s drifting grating.

### Discrete Cl^-^ hotspots at the onset of GABA responses

At the onset of inhibitory activity, the spatiotemporal dynamics of Cl^−^ are expected to follow Brownian diffusion from a punctate source (the synapse) along the dendrite (see Material and Methods and Supp. Fig. 1). In a set of experiments, fluorescence was acquired at 20 Hz to maximise temporal resolution and measured along a selected dendrite (Fig. 3a). Fluorescence was corrected for offset and photobleaching by acquiring the mCRISPRed signal at the beginning and end of each sequence; given the modest degree of photobleaching, the correction was approximated with a linear function. To detect the onset of Cl^−^ transients and suppress slow baseline drift, the temporal derivative of the signal was computed as:

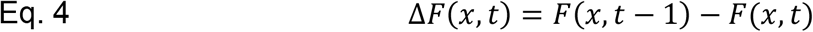

where x is the coordinate along the dendrite. This operation yields a positive value when [Cl^−^]i increases at position x between consecutive frames.

The matrix *ΔF(x, t)* was filtered along x by convolution with a Gaussian kernel to account for the spatial blurring introduced by Cl^−^ diffusion during the camera integration period (see also Supp. Fig. 1). The matrix was then Z-scored against baseline statistics from the pre-stimulus period and thresholded at Z=3, corresponding to p < 0.002, to extract significant Cl^−^ transients (Fig. 3b, middle panel). Given that a large number of pixels were tested simultaneously, a conservative threshold was chosen. All non-zero elements of the Z-score matrix were considered valid Cl^−^ transients. Their distribution along the dendrite was obtained by counting valid transients across the entire duration of the response (Fig. 3b, bottom panel), revealing clustering of transients in discrete hotspots.

To estimate the spatial extent of these active regions, the distribution was modelled with a Gaussian mixture, using the Bayesian information criterion (BIC) to identify the optimal number of Gaussian components while avoiding overfitting (Fig. 3b, bottom panel). To assess whether the observed clustering could arise from the random superposition of events, we randomised the positions of Cl^−^ transients for each dendrite while preserving their size, number, and the total dendrite length, and compared the nearest-neighbour statistics of the randomised data with those of the observed data. Figure 3c shows that observed transients were grouped significantly closer together than expected by chance (two-sided Kolmogorov-Smirnov test, p<<0.001), across 28 dendrites imaged in 3 separate acquisition rounds. The distribution of hotspot sizes, expressed as the full width at half maximum of the Gaussian model, is shown in Fig. 3d.

If hotspots reflect stable sites of Cl^-^ influx, rather than stochastic events, they should recur at consistent dendritic locations across repeated stimulus presentations. To test this, each dendrite was imaged across 2 or 3 trials and the density profiles of Cl^−^ hotspots were compared by computing their cross-correlation. Averaged across all imaged dendrites (n=25), the cross-correlation shows a clear central peak at lag zero, indicating that hotspots tend to recur at the same dendritic locations across trials (Fig. 3e,f). This spatial reproducibility disappeared after random circular permutation of the density profiles, confirming that the overlap is not due to chance.

### *In vivo* imaging of GABAergic Cl^-^ fluxes after visual stimulation

We demonstrated the photostability of iClima and its suitability for extended *in vivo* sessions by the epileptic seizure recordings described in Supp. Fig. 3. In principle, the high Cl^−^ affinity of iClima should detect small chloride fluxes driven by physiological inhibitory activity. To demonstrate this capability *in vivo*, we monitored [Cl^−^]i in pyramidal neurons of the mouse visual cortex during visual stimulation. After ten days of habituation to head restraint and the imaging setup, four iClima-expressing mice were imaged across multiple sessions using two-photon microscopy in the awake head-restrained condition. iClima was excited at the isosbestic wavelength (830 nm) and responses were computed as the change in g/r ratio relative to baseline. Baseline periods consisted of a uniform gray screen followed by a 5 s presentation of a drifting grating at 95% Michelson contrast, with mean luminance matched to the gray baseline (Fig. 4a,b). In Supp. Fig. 4, we verified that the luminance changes of the monitor did not interfere with the detection of Cl^-^ transients. Response amplitudes were estimated by fitting individual traces with a logistic function for the rising phase, as described below. Visual stimulation induced robust and reproducible Cl^−^ transients, with an average peak of 3.6% (n=366 neurons, 4 mice). To our knowledge, this is the first optical detection of Cl^−^ transients at single-cell resolution evoked by physiological sensory stimulation *in vivo* in the mouse cortex.

As discussed above, the pH robustness of iClima makes any contribution from intracellular pH fluctuations unlikely. However, the observed signal could in principle reflect haemodynamic responses of brain tissue that differentially affect transmission in the red and green channels. To exclude both artefacts simultaneously, we repeated the same experiment using LSSmClopHensor, which has much greater pH sensitivity than iClima and similar emission spectra, making it equally susceptible to any haemodynamic effect. If the observed transients were attributable to pH changes or haemodynamic responses, they should be detected, or even amplified, by LSSmClopHensor. We imaged 714 neurons in 7 mice expressing LSSmClopHensor at 910 nm, its isosbestic wavelength. As shown in Figure 4c,d, even after averaging across multiple trials, neurons, and mice, the response was barely detectable. This result demonstrates: i) the Cl^−^ transients detected by iClima are not caused by pH shifts, since the highly pH-sensitive LSSmClopHensor displays a smaller response than the pH-robust iClima; ii) the signals are not attributable to haemodynamic artefacts; and iii) iClima is substantially more sensitive than LSSmClopHensor for detecting physiological Cl^−^ transients *in vivo*.

To exclude potential confounds arising from locomotion and changes in behavioural state, subsequent experiments were performed in isoflurane-anaesthetised mice, allowing more controlled assessment of stimulus-evoked Cl^−^ responses. Four iClima-expressing mice were imaged across multiple sessions; a total of 1662 neurons were recorded across all stimulus conditions, of which those exhibiting a significant response to each stimulus were included in the respective analyses. The primary stimulus was a full-field luminance step: following a dark baseline, the display was switched to a bright uniform screen for 20 s, then returned to darkness. This paradigm is known to evoke strong excitatory and inhibitory responses in V1 neurons, with inhibitory components peaking shortly after stimulus onset^17^.

Using iClima, we observed clear Cl^−^ transients in the soma of pyramidal neurons at stimulus onset, with an average peak of 3.3% (n=817 neurons, Fig. 4e). We additionally presented drifting horizontal gratings of two different durations. A Cl^−^ increase was detected in both conditions, with average peaks of 4.9% and 5.1% for the 5 s and 10 s gratings respectively, suggesting that the Cl^−^ response saturates rapidly and is largely independent of stimulus duration within this range (Fig. 4f,g).

The shape of the Cl^−^ transient carries information about the kinetics of inhibitory input and Cl^−^ extrusion that complements the amplitude measurements. The rising phase, driven by GABA_A_-mediated Cl^−^ influx, is well described by a logistic function; the post-stimulus recovery, reflecting KCC2-mediated extrusion, follows a single exponential decay. We therefore fitted the response to the 5 s grating with a piecewise model consisting of a logistic growth joined at the peak to an exponential decay. This model provides an excellent description of the observed responses across all four mice, as demonstrated by the fits and residual analysis (Fig. 4f and Table S1). This piecewise model is used in the analysis of Figure 5 to isolate the Ca^2+^ signal from the mixed Ca^2+^/Cl^−^ response at 920 nm.

**Figure 5.**
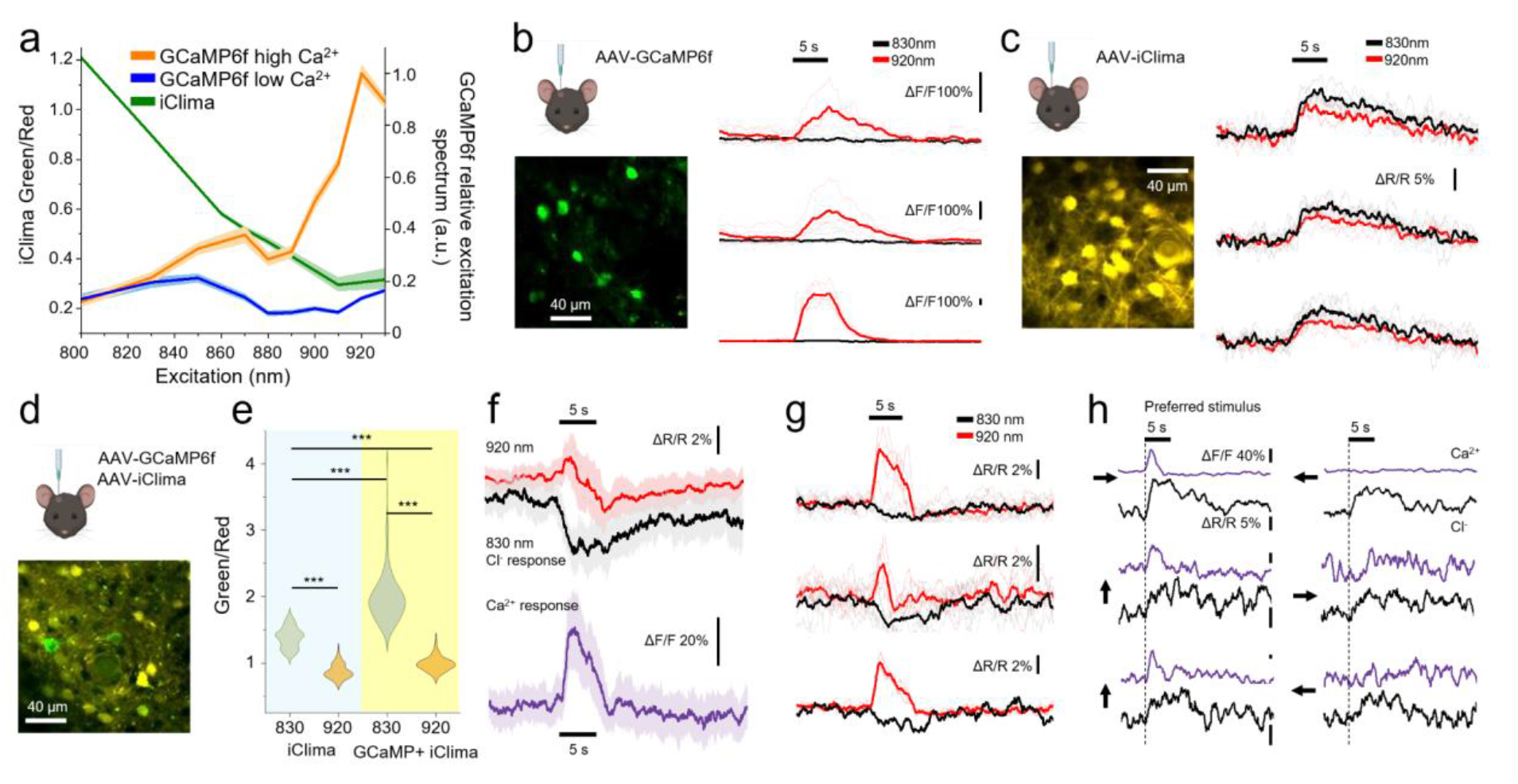
Simultaneous Cl^−^ and Ca^2+^ imaging. **a**) Comparison of two-photon excitation spectra of GCaMP6f (Ca^2+^-bound and Ca^2+^-unbound states) and iClima. Fluorescence was normalised to the squared photon number delivered to the sample. **b**) Representative field of view (920 nm) and responses to visual stimulation from 3 neurons in mice expressing GCaMP6f only. Responses are present at 920 nm but absent at 830 nm, confirming Ca^2+^- independence of the iClima signal at the isosbestic wavelength. **c**) Representative field of view (830 nm) and responses from 3 neurons in mice expressing iClima only, expressed as *ΔR*/*R*_*baseline*_ of the red/green ratio. **d**) Representative field of view (920 nm) in a mouse co-expressing both sensors. **e**) Distribution of resting g/r ratios in iClima-only (n=223) and co-expressing neurons (n=285). The higher ratio in co-expressing cells reflects GCaMP6f emission in the green channel (two sided Mann-Whitney test, ^***^ p < 0.001). **f**) Median ΔR/R traces (n=104 co-expressing neurons) at 920 nm (red) and 830 nm (black), and the isolated Ca^2+^ response after Cl^−^ component subtraction (lower panel). Shaded areas: median absolute deviation. **g**) Single-cell fluorescence dynamics from 3 representative co-expressing neurons. Cl^−^ transients appear as downward deflections; Ca^2+^ responses as upward deflections. Thin lines: individual trials; bold lines: trial averages. **h**) Ca^2+^ (purple) and Cl^−^ (black) responses from neurons imaged during drifting gratings of different orientations. Each row shows a neuron responsive (left) or non-responsive (right) to a specific direction (black bar, 5 s). Lines represent trial averages. Horizontal bars indicate stimulus timing in all panels. Illustrations in panels b, c and d generated with BioRender.com.

### Simultaneous optical measurement of inhibitory drive and excitatory output in single neurons

Two-photon calcium imaging with GCaMP indicators has become the dominant approach for monitoring neuronal activity in the living brain. However, this capture the overall excitation state resulting by synaptic integration, remaining entirely blind to the inhibitory component of synaptic integration. iClima provides an optical readout of inhibitory synaptic drive through Cl^−^ fluxes. Crucially, the spectral properties of iClima make it compatible with simultaneous Ca^2+^ imaging using GCaMP6f, enabling the concurrent measurement of Cl^-^ current and excitatory output within the same identified neuron *in vivo*.

The basis for this multiplexing strategy lies in the two-photon excitation spectra of the two sensors. At the iClima isosbestic wavelength (830 nm), GCaMP6f fluorescence is Ca^2+^-independent, as confirmed by the absence of visually evoked Ca^2+^ responses at this wavelength (Fig. 5a,b, Supp. Fig. 5). Consequently, the g/r fluorescence ratio obtained at 830 nm provides a selective readout of Cl^−^ transients, free from any Ca^2+^ contribution. At 920 nm, by contrast, the fluorescence is modulated by both Ca^2+^ and Cl^−^. Since Cl^−^-dependent quenching of iClima arises from static quenching, the Cl^−^- dependent modulation of fluorescence at 920 nm mirrors that at 830 nm in both kinetics and waveform, differing only in a small amplitude scaling factor (Fig. 5c). This property makes it possible to isolate the Ca^2+^ component at 920 nm by subtracting a noiseless analytical model of the Cl^−^ response, derived from the 830 nm channel, on a cell-by-cell basis, avoiding the noise amplification that would result from direct subtraction of two raw imaging channels (see Supplementary Information and Supp. Fig. 6). The Cl^−^ response model used here is the same piecewise logistic plus exponential model introduced above, which provides an adequate characterisation of the response to the 5 s grating stimulus used in this experiment. To implement this approach *in vivo*, mice were co-injected with AAV vectors driving Cre-dependent expression of iClima and GCaMP6f, together with an AAV expressing Cre recombinase under the CaMKII promoter, thereby restricting dual expression to excitatory pyramidal neurons.

Figure 5d shows a representative field of the visual cortex imaged at 10 Hz in spiral scan mode at 920 nm, illustrating the heterogeneity in dual sensor expression levels across neurons. The presence of Ca^2+^-unbound GCaMP6f contributes additional fluorescence to the green channel in co-expressing neurons, shifting their resting g/r ratio upward relative to iClima-only cells — a shift confirmed statistically across the imaged population (Fig. 5e; two-sided Mann-Whitney test, p<0.001) and used to identify co-expressing neurons for subsequent analysis. In co-expressing neurons (n=104 neurons, 3 mice), visual stimulation with drifting gratings produced clearly distinguishable Ca^2+^ and Cl^−^ transients (Fig. 5f,g). By convention, with the g/r ratio as the readout, Ca^2+^ responses appear as positive deflections and Cl^−^ increases as negative deflections, as they have opposite effects on green fluorescence. At 920 nm, a biphasic dynamic is observed: a fast upward transient reflecting the Ca^2+^ response, followed by a slower downward deflection reflecting the Cl^−^ transient. At 830 nm, only the downward Cl^−^ component is present, confirming Ca^2+^- independence at this wavelength (Fig. 5f). After subtraction of the scaled Cl^−^ model from the 920 nm signal, the isolated Ca^2+^ response is recovered as shown in the lower panel of Fig. 5f. Importantly, Ca^2+^ and Cl^−^ responses can be resolved in individual neurons on a single-trial basis (Fig. 5g), establishing the practical utility of the approach for trial-by-trial analyses.

Figure 5h shows responses of three neurons to their preferred and opposite stimuli; arrows indicate the drift direction of the grating. Inspection of the Ca^2+^ responses reveals that the upper neuron is direction-selective, while the other two are orientation-selective. Remarkably, in this limited sample, the amplitude of the Cl^−^ transient are independent of the stimulus direction or orientation, consistent with the idea that stimulus specificity in cortical neurons is primarily conferred by excitatory inputs.

## Discussion

iClima advances genetically encoded Cl^-^ imaging by combining high Cl^−^ affinity, pH robustness under physiological conditions and photostability sufficient for prolonged in vivo two-photon imaging, a set of properties not previously achieved in a single indicator ^18,19^. The K_d_ of 3.5 mM at pH 7.2 places iClima firmly within the physiological range of [Cl^−^]i in mature cortical neurons (5–10 mM), representing a substantial improvement over LSSmClopHensor and SuperClomeleon^11^, whose FRET-based design presents additional technical challenges for *in vivo* implementation. Equally important is iClima pH insensitivity conferred by the high pK_a_ of 7.84. Since neuronal intracellular pH sits well below this value (7.1–7.2), iClima is predominantly protonated under resting conditions, and is minimally affected by the small pH fluctuations that accompany physiological neuronal activity. We have demonstrated that an undetected pH change of 0.1 units — well within the range expected during normal inhibitory activity — introduces less than 4% error in the Cl^−^ estimate with iClima, compared to approximately 20% with LSSmClopHensor. This property is not merely a technical convenience but it has profound consequences: it is what makes single-wavelength dynamic imaging feasible, removing the need for simultaneous pH correction and dramatically simplifying the experimental design for *in vivo* applications, including fiber photometry. Alongside the substantially improved photostability of both iClima components relative to their LSSmClopHensor counterparts, these properties collectively enable imaging paradigms that were previously out of reach.

The most immediate demonstration of iClima’s capabilities is the first optical detection of Cl^−^ transients evoked by physiological sensory stimulation *in vivo*. Previous *in vivo* Cl^−^ imaging in the mouse cortex with single-cell resolution has been confined either to static measurements of baseline [Cl^−^]i and its slow homeostatic modulation^4,5,9^, or to dynamic measurements under strongly non-physiological conditions such as seizures and hypercapnia^4^. Few studies attempted to measure Cl^-^ transients in response to physiological activity by either fiber photometry^20–22^ or by two photon imaging in the olfactory system of drosophila^23^. Both studies do not have a clear single neuron resolution and are potentially affected by the complex dependency of SuperClomeleon and Clomeleon on intracellular pH. Here we show that it is possible to resolve sensory-evoked Cl^−^ transients of 3–5% in individual neurons of awake, behaving mice thus opening an entirely new experimental window onto inhibitory circuit function *in vivo*. The use of LSSmClopHensor as an internal control is a powerful validation, exploiting its greater pH sensitivity and similar spectral properties to rule out pH artefacts and haemodynamic contributions simultaneously. The near-absence of detectable responses with LSSmClopHensor under identical stimulation conditions provides compelling evidence that the iClima signals reflect genuine Cl^−^ dynamics. The subcellular hotspots analysis further demonstrates that iClima has the sensitivity and photostability to resolve spatially discrete Cl^−^ hotspots along individual dendrites and in perspective, these studies could be replicated *in vivo* where the heterogeneity of the localization and operation of inhibitory synapses is maximal.

Perhaps the most significant new capability introduced here is the simultaneous imaging of Ca^2+^ and Cl^−^ transients in the same identified neuron *in vivo*. Two-photon Ca^2+^ imaging with GCaMP indicators has transformed systems neuroscience by making it possible to monitor the activity of hundreds of neurons simultaneously in the living brain. However, Ca^2+^ signals report neuronal activity, leaving the underlying process of integration of excitatory and inhibitory inputs entirely invisible. iClima contributes to fill this gap by providing a spectrally compatible optical readout of inhibitory Cl^-^ currents that can be acquired simultaneously with GCaMP6f using a simple two-wavelength switching protocol. A key result is the demonstration that Ca^2+^ and Cl^−^ responses can be resolved on a single-trial basis in individual neurons *in vivo* establishing that the approach is practical for the kind of trial-by-trial analyses that are central to modern systems neuroscience.

Our proof-of-principle observations suggesting that in orientation selective cells the amplitude of Cl^−^ transients are independent of stimulus orientation and direction, is consistent with the hypothesis that stimulus specificity in cortical pyramidal neurons is primarily sculpted by the tuning of their excitatory inputs rather than by matched inhibition^24–26^. This is a contentious issue in the field: while several studies using Ca^2+^ imaging and optogenetic manipulations of identified interneuron subtypes have concluded that inhibitory neurons, particularly parvalbumin-expressing cells, are broadly tuned and act primarily as a gain control mechanism, others have reported interneuron tuning comparable to that of excitatory cells, leading to competing models of inhibitory circuit function in visual cortex^27–29^. Our somatic Cl^−^ measurements are likely dominated by perisomatic inhibition from parvalbumin-expressing basket cells, which are known to be broadly tuned^24,26^ and therefore we do not exclude the possibility that dendritic inhibition, delivered by somatostatin-expressing interneurons, may exhibit a degree of stimulus selectivity that is not captured by somatic measurements. Resolving this question will require the kind of compartment-specific Cl^−^ imaging that iClima’s spatial resolution and sensitivity are uniquely positioned to enable, and is a direction we are actively pursuing.

A current limitation of the combined Ca^2+^/Cl^−^ imaging protocol demonstrated here is that the two excitation wavelengths (830 nm and 920 nm) are acquired in successive trials rather than simultaneously, requiring the model-based subtraction approach described in the Methods to separate the two signals. While this strategy is effective, inter-trial variability in neuronal responses inevitably limits the precision of the signal separation on a trial-by-trial basis. Two technical strategies could overcome this limitation in future implementations. The simpler approach is frame-interleaved dual-laser acquisition, in which two Ti:Sapphire lasers are alternated between 830 nm and 920 nm on a frame-by-frame basis using fast electro-optical shutters, reducing the effective frame rate per channel by half. This solution is immediately implementable on existing dual-laser two-photon systems and, when combined with resonant scanning to maintain adequate temporal resolution, represents a practical near-term upgrade. A more powerful solution is pulse-interleaved excitation combined with high-speed time-resolved detection. In this approach, the pulses of two phase-locked lasers are interleaved at the nanosecond timescale — exploiting the ∼12.5 ns interpulse interval of a standard 80 MHz Ti:Sapphire laser to accommodate pulses from a second laser offset by ∼ 6 ns — so that fluorescence from each excitation wavelength can be assigned unambiguously based on photon arrival time using time-correlated single photon counting (TCSPC) electronics or high-speed analogue digitisers^30,31^. This achieves genuine simultaneous acquisition with no reduction in frame rate. Looking forward, iClima is well positioned to address open questions about how the the interplay between excitation and inhibition evolves during development, across brain states, or in disease models where dysregulation of the inhibitory feedback plays a central role, including epilepsy, autism spectrum disorders, and schizophrenia — contexts in which a tool capable of directly and simultaneously monitoring both sides of the balance at single-cell resolution offers capabilities that have not previously existed.

## Acknowledgements

GMR is supported by Telethon Grant GMR24T1119, and by Fondation Jérome Lejeune project No.2501-2025A. AF and GMR are supported by PRIN 2022 2022A58M7F_LS5. AF is supported by Telethon Grant GMR24T1085. CL is supported by Telethon Grant GMR23T1055. RSE is supported by UKRI Future Leaders Fellowship MR/Y017552/1. JSO is supported by the Medical Research Council (MC_UP_1201/4) as part of United Kingdom Research and Innovation. Schematic illustrations were created with BioRender (BioRender.com).

## Materials and Methods

### Generation and selection of mClover3 mutations

Mutations conferring elevated proton affinity and enhanced Cl^−^ sensitivity were identified by directed evolution of E^2^GFP. Briefly, random mutagenesis libraries were generated by error-prone PCR and screened in a high-throughput format for fluorescence brightness, *pK*_*a*_, and Cl^−^ dissociation constant (*K*_*d*_). An intermediate variant with substantially elevated *pK*_*a*_ served as the template for site-saturation mutagenesis at positions identified as key determinants of proton and anion affinity. Saturation mutagenesis was performed using the OmniChange method^32^, which enables simultaneous saturation of multiple codons in a single assembly reaction. Candidate mutations were ranked by the lower bound of the 94% highest-density interval of the Bayesian *pK*_*a*_ posterior, providing a conservative selection criterion robust to measurement noise. The identified substitutions were introduced into the mClover3 backbone to generate mClover3*.

The green module (mClover3*) was generated on an mClover3 template and differs from it at eleven positions (avGFP numbering throughout). Seven substitutions rebuild the E^2^GFP/superfolder background required for anion sensing and folding: F64L and G65T restore the E^2^GFP chromophore; H203Y installs the Tyr203 that forms the halide-binding site of E^2^GFP; A69Q and Y149N revert the Clover-lineage residues to those retained in avGFP, E^2^GFP and sfGFP, Gln69 being a constituent of the anion-binding pocket; and K206V replaces the monomerising lysine of mClover3 (A206K) with the superfolder valine (A206V). C160S eliminates the unpaired cysteine introduced by mClover3 at position 160, which is glycine in avGFP, EGFP, E^2^GFP, Clover, sfGFP and mGreenLantern. The protein further carries the superfolder set S30R, Y39N, F99S, N105T, Y145F, M153T, V163A and I171V, common to sfGFP and the Clover lineage. The remaining four substitutions, D21H, A87T, N185T and S202N (HTTN), were introduced to raise the apparent *pK*_*a*_ to 8.

For mammalian expression, the coding sequences of mClover3* and mCRISPRed were codon-optimised for *Mus musculus* using DNAchisel^33^. The final iClima insert (mClover3*- HTTN–linker–mCRISPRed) was cloned by ligation into the pCAG expression vector at EcoRI and NotI restriction sites, placing the construct under the CAG promoter. The resulting expression cassette was verified by Sanger sequencing prior to functional characterisation.

### Cell culture

Calibration experiments were carried out in murine embryonic fibroblasts (MEF; American Type Culture Collection), U2-OS cells, and mouse primary hippocampal neurons. MEF and U2-OS cells were maintained in Dulbecco’s Modified Eagle Medium (DMEM; Invitrogen, Carlsbad, CA, USA) supplemented with 10% fetal bovine serum (FBS), 4 mM L-glutamine, 1 mM sodium pyruvate, 100 U/mL penicillin, and 100 µg/mL streptomycin (Invitrogen). Cells were maintained at 37 °C in a humidified incubator with 5% CO_2_.

Mouse primary hippocampal neurons were prepared from C57B6J mouse embryos at E17– 18 (approval protocol: 22418.N.ZBX). Hippocampi were dissected under a stereomicroscope in ice-cold Ca^2+^/Mg^2+^-free HBSS. Tissue was incubated for 10–15 min in 0.25% trypsin (Thermo Fisher Scientific) at 37 °C. The trypsin solution was then removed and replaced with Neurobasal medium (Thermo Fisher Scientific) supplemented with 10% FBS (Thermo Fisher Scientific), 1% GlutaMAX (Thermo Fisher Scientific), and 1% penicillin/streptomycin (PenStrep, Thermo Fisher Scientific). The digested tissue was mechanically dissociated, cells were counted and plated on 25 mm coverslips or petri dishes coated with poly-L-lysine (0.1 mg/mL) at a density of 200 cells/mm^2^. Two hours after plating, the medium was replaced with Neurobasal medium supplemented with 1% B27 (Thermo Fisher Scientific), 1% GlutaMAX, and 1% PenStrep. Cultures were maintained in a humidified incubator at 37°C, 5% CO_2_ until at least 14 days *in vitro*.

### Microscopy

For transfection cells were trypsinised, pelleted by centrifugation, and washed with PBS. Cells were resuspended in the appropriate electroporation buffer containing plasmids encoding mClover3*, mCRISPRed, or iClima. Electroporation was performed using the NEON™ Electroporation System (Invitrogen) with a single square-wave pulse (1350 V, 30 ms), according to the manufacturer’s instructions. Following electroporation, cells were seeded onto 35 mm dishes, allowed to recover for 48 hours, and subsequently subjected to experimental treatments prior to imaging. U2-OS cells stably expressing the iClima construct were used without further transfection.

Two-photon imaging was performed on a Bruker Ultima microscope equipped with a Chameleon Ultra II Ti:Sapphire laser (Coherent) at eight excitation wavelengths as shown in Fig 1d. Emitted fluorescence was detected through green and red band-pass filters (527/70 nm and 607/70 nm respectively; BrightLine). Imaging was conducted at 34 °C using a 20× water immersion objective (NA 1.05; Olympus).

One-photon characterisation of iClima was conducted on two confocal microscopes. A Leica Stellaris confocal microscope equipped with a white-light laser was used to acquire emission spectra between 465 and 695 nm (step size 10 nm, bandwidth 30 nm) at excitation wavelengths of 440, 445, and 450 nm, and then every 10 nm up to 490 nm. Emission spectra were integrated over two wavelength ranges corresponding to the green (520–555 nm) and red (590–625 nm) channels. A Leica SP8 confocal microscope was used for imaging at 442 nm and 488 nm excitation; emitted fluorescence was detected through green and red band-pass filters (500–545 nm and 570–700 nm respectively). Characterisation of iClima in neurons *in vitro* is described in the Live Imaging section below.

### pH and chloride calibration

For pH calibration, MEF cells were incubated with calibration buffer (20 mM HEPES, 0.6 mM MgSO_4_, 38 mM sodium gluconate, 100 mM potassium gluconate, zero Cl^−^) supplemented with an ionophore mixture to equilibrate intracellular and extracellular ion concentrations. The ionophore mixture contained: 10 µM tributyltin chloride (Cl^−^/OH^−^ exchanger), 5 µM nigericin (K^+^/H^+^ exchanger), 5 µM valinomycin (K^+^ ionophore), and 5 µM carbonyl cyanide p-chlorophenylhydrazone (CCCP; protonophore). Cells were washed six times for three minutes with ionophore solution to allow complete ion equilibration. pH was varied across the range 6 to 8 by adjusting the extracellular buffer composition. For chloride calibration, cells were incubated with an ionophore mixture at fixed pH 7.2 with chloride concentrations ranging from 0 to 138 mM, adjusted by substituting sodium gluconate with NaCl.

Images were processed using custom MatLab scripts. The ROI based green and red fluorescence of each cell was corrected for background offset and bleed-through as described above. For pH calibration, the g/r ratio at each excitation wavelength was decomposed as *X*_*pH*_ = *δ · X*_*pH=6*_+ *ε X*_*pH=8*_. pK_a_, and the pK_a_ was determined by fitting *θ* to Eq. 1. An analogous procedure was applied for chloride calibration, in which the g/r ratio was fitted with Eq. 2 to obtain the Kd.

### Live-imaging experiments in primary neurons

#### Neuronal transfection

Primary hippocampal neurons were transfected at DIV 14 using Lipofectamine 2000 (Thermo Fisher Scientific) according to the manufacturer’s protocol, using 0.5 µg cDNA per 25 mm coverslip. Conditioned medium was replaced after transfection and neurons were imaged at DIV 17–20.

#### Live imaging of Cl^-^ transients in cultured neurons

For live imaging, coverslips were transferred to an imaging chamber (∼200 µL volume; Quick Exchange Platform, Warner Instruments) containing Tyrode’s solution (140 mM NaCl, 3 mM KCl, 2 mM CaCl_2_, 1 mM MgCl_2_, 10 mM HEPES pH 7.4, 10 mM glucose) supplemented with 25 µM CNQX and 50 µM D-APV (D-2-amino-5-phosphonovaleric acid) to suppress glutamatergic transmission. Imaging was performed at 30 °C. Field stimulation was delivered through platinum–iridium electrodes using an AM2100 stimulator (A-M Systems) with 1 ms current pulses at 20 Hz. Baseline fluorescence was acquired prior to stimulation. For pharmacological experiments, bicuculline (30 µM, Tocris) was bath-applied, and KCC2 was inhibited by application of VU0463271 (0.5 µM, Tocris).

Fluorescence imaging was performed on an inverted wide-field epifluorescence microscope (Olympus IX81) using a 40× objective (Fig. 2) or 60× objective (Fig. 3) oil-immersion objective (NA 1.35). Illumination was provided by a broad-spectrum LED source (pE-300ultra, CooLED, Andover). The optical configuration for iClima consisted of a 435–455 nm excitation filter, a 495 nm dichroic mirror, a 500–550 nm emission filter (green channel), and a 565–615 nm emission filter (red channel). Rapid alternation between emission channels was achieved using an Olympus U-FFWO fast filter wheel. Images were acquired with an Orca C4742-80-12AG CCD camera (Hamamatsu Photonics).

#### Image analysis

Image sequences were processed in MatLab using a semi-automated pipeline. ROIs corresponding to individual neuronal somata or dendritic segments were drawn manually on the mean fluorescence image. After motion correction, the mean green and red fluorescence was extracted for each ROI across all frames. Signals were first corrected for background offset, measured from a cell-free region of each image, and subsequently corrected for bleed-through (see Supplementary Information). No temporal filtering was applied. The corrected signals were used to compute the green-to-red fluorescence ratio (r/g) across the entire recording. The mean ratio during the baseline period was defined as R_*baseline*_, and the normalised change in fluorescence ratio was calculated as defined in Eq. 3.

#### Analysis of Cl^-^ hotspots

The green iClima signal from cultured neurons was acquired at 20 Hz with a camera integration time of 10 ms per frame. Photobleaching was assessed by acquiring a reference red frame at the beginning and end of each recording and was corrected by fitting a linear function to these two measurements, approximating the bleaching time course as a linear ramp. The frame-by-frame variation matrix of the green signal was computed as defined in Eq. 4.

At any given time point, the signal measured along the dendrite reflects both the local Cl^−^ concentration and noise. The Cl^−^-dependent component is spatially correlated along x (the coordinate along the dendrite), since Cl^−^ ions diffuse during the camera integration period. Noise, by contrast, is spatially uncorrelated between adjacent pixels. In the first analysis step, uncorrelated noise was suppressed while preserving the spatially correlated Cl^−^ signal by convolving the signal at each time point along the x dimension with a Gaussian kernel. The kernel width was chosen to match the spatial resolution limit imposed by Cl^−^ diffusion during the integration period, as derived below.

The diffusion coefficient of Cl^−^ ions in water, *D*_*Cl*_, can be estimated from the Nernst-Einstein equation linking ionic diffusivity to solution conductivity, yielding *D*_*Cl*_ =2.03 × 10^3^ µm^2^/s. In the intracellular environment, diffusion is substantially reduced by molecular crowding and viscosity; we therefore assumed *D*_*Cl*_ = 1.0 × 10^3^ µm^2^/s as a conservative estimate (see Kuner & Augustine 2000^34^). If at time zero a point-like Cl^−^ source is activated at a synapse and ions begin diffusing, the expected concentration distribution at time *t* and distance *x* from the source is:

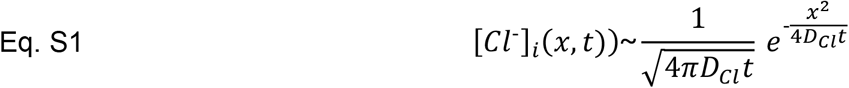

This expression shows that Cl^−^ disperses following a Gaussian profile with standard deviation:

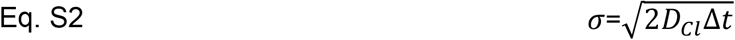

which can be interpreted as the mean distance reached by Cl^−^ ions from the source after time Δt (Supp. Fig. 1a). The camera integration time is Δt=10 ms. Since the precise timing of receptor opening within the integration window is unknown, we conservatively assume that, on average, the data reflect a diffusion period equal to half the integration time, i.e. Δt=5 ms. Substituting into Eq. S2 yields an expected diffusion length of σ ≈3 µm. This means that the observed Cl^−^ distribution is effectively low-pass filtered by the diffusion process, with a characteristic spatial scale of approximately 3 µm (shaded area of Supp. Fig. 1a).

To maximise the signal-to-noise ratio while preserving the spatial structure of the Cl^−^ signal, we applied a matched filter: the signal matrix was convolved along the spatial dimension with a Gaussian kernel whose standard deviation was set to σ =1.5 µm, matching the spatial autocorrelation length of the diffusion-blurred Cl^−^ signal. This approach is optimal in the sense that it maximises the detectability of spatially correlated Cl^−^ transients relative to uncorrelated pixel noise, without introducing major spatial smoothing beyond the scale already imposed by diffusion. Note that this filtering does not artificially broaden the detected Cl^−^ hotspots beyond their diffusion-limited extent, since the kernel width is smaller than the characteristic diffusion length σ ≈ 3 µm. Supp. Fig. 1b,c shows the ΔF map before and after Gaussian filtering. After filtering, the map was Z-scored as:

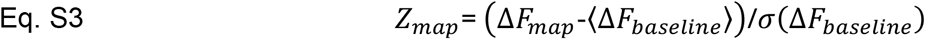

where *ΔF*_*map*_ is the filtered raster map, ⟨*Δ*F_*baseline*_⟩ is the mean value of the filtered raster map computed at each dendrite location during the baseline period (2 s, 60 frames), and *σ*(*ΔF*_*baseline*_) is the corresponding standard deviation. After filtering, the map was Z-scored against baseline statistics (Eq. S3; baseline period: 2 s, 60 frames) and thresholded at Z>3 (p<0.002, Supp. Fig. 1). Contiguous pixels along x within each frame were classified as a single Cl^−^ transient, characterised by its spatial extent and position along the dendrite. To assess whether the observed clustering exceeded chance, surrogate datasets were generated by randomising transient positions along each dendrite while preserving their number, size, and total dendrite length; observed and surrogate nearest-neighbour distance distributions were compared using the Kolmogorov-Smirnov test.

**Supplementary Figure 1.**
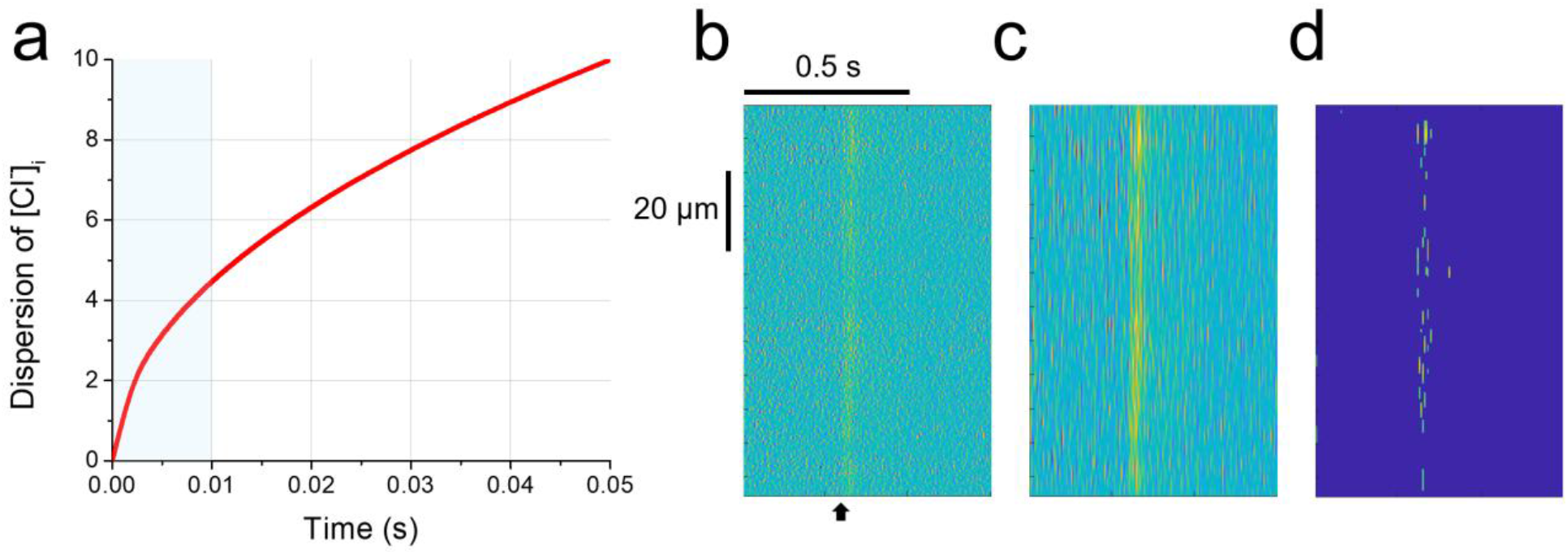
Identification of Cl^-^ hotspots. **a**) Mean diffusion distance of Cl ion from the point of entry as a function of time. The shaded area indicates the integration time of the camera. **b**) Raw raster plot of *ΔF*_*map*_. The arrow head indicates the timing of the onset of the stimulus. **c**) raster map after convolution with the Gaussian filter. **d)** Raster map after Z-scoring and thresholded for Z>3.

### *In vivo* imaging

#### Viral injections and chronic implants

*In vivo* expression of iClima, LSSmClopHensor, and GCaMP6f was achieved through AAV-mediated transduction. Expression was restricted to CaMKII-expressing excitatory neurons using a Cre-dependent strategy, in which AAV vectors carrying floxed (DIO) sensor genes (AAV9.EF1α.DIO(LSSmClopHensor), AAV9.EF1α.DIO(iClima), AAV9.CAG.DIO(GCaMP6f)) were co-injected with AAV9.CaMKII.Cre. The titres of the sensor-expressing viral vectors were approximately 8.7 × 10^11^ vg/mL; the titre of the Cre-expressing virus was 2.1 × 10^11^ vg/mL.

Mice (C57BL/6J, 3-4 months) were anaesthetised by intraperitoneal injection of ketamine/xylazine (100 mg/kg and 5 mg/kg body weight respectively). Animal of both sexes were used following approval protocols 129/2000-A and 1060/20.Two injections of 500 nL each were made at a depth of 300 µm in the visual cortex equally spaced around stereotaxic coordinates AP = 0 mm and ML = 2.5–3.5 mm from lambda. Imaging was performed starting 3 weeks after viral injection to allow sufficient sensor expression.

A 3 mm craniotomy was performed over V1. The craniotomy was covered with a chronic cranial window consisting of two circular glass coverslips (inner diameter 3 mm, outer diameter 5 mm; 0.16 mm thickness) bonded with optical glue. A custom steel head bar (<1 g) was fixed to the skull caudal to the craniotomy using Metabond dental cement (Parkell), followed by a layer of Paladur (Kulzer).

#### Preparation of the mouse for in vivo imaging

For awake imaging, mice were habituated to head fixation over 9 sessions across ten days prior to imaging. Mice were positioned under the two-photon microscope at a distance of 30 cm from the stimulus monitor, with the eye contralateral to the imaged hemisphere facing the screen. Care was taken to shield the photomultiplier tubes from monitor light. Imaging was performed at depths of 50–300 µm from the brain surface, corresponding approximately to cortical layers 1–3, at 2 frames per second. Unless otherwise specified, animals were maintained under isoflurane anaesthesia (1–2% in oxygen), with the concentration adjusted to maintain a stable physiological state.

#### Visual stimulations

Visual stimuli were generated using a custom MATLAB application employing Psychophysics toolbox and displayed on a LCD monitor (1920x1080 pixels, refresh rate 75Hz, size 54x31 cm) positioned 30 cm from the animal. The monitor was gamma-corrected and luminance was calibrated using a Minolta luminometer at several luminance points to correct for nonlinearity.

Synchronisation between visual stimuli and imaging acquisition was achieved using a phototransistor monitoring a concealed corner of the monitor screen, which generated TTL pulses recorded at 5 kHz together with the microscope frame trigger, providing precise temporal alignment of stimuli with imaging frames and electrophysiological recordings.

Two categories of visual stimuli were used. The first was a full-field luminance step, in which monitor luminance was set to ∼0 cd/m^2^ during a 30 s baseline period, stepped to 10 cd/m^2^ for 20 s, and then returned to ∼0 cd/m^2^ during a 20 s recovery period. The second category consisted of full-field drifting grating stimuli. For gratings, the baseline consisted of a uniform gray screen at a mean luminance of 5 cd/m^2^ for 30 s. Square-wave gratings (spatial frequency: 0.02 cycles/degree; temporal frequency: 2 Hz) were presented for either 5 s or 10 s depending on the experimental protocol, and consisted of alternating white (10 cd/m^2^) and black (∼0 cd/m^2^) bars, yielding 95% Michelson contrast ((Lmax − Lmin)/(Lmax + Lmin)). For direction selectivity experiments, gratings were presented at 8 orientations separated by 45 degrees; 0° corresponded to motion in the tail-to-head direction, with other orientations defined accordingly. Each condition was repeated for several trials with an inter-trial interval of 30 s.

#### In vivo seizure

A 4 mm craniotomy was performed over V1 and sealed with a 5 mm glass coverslip secured to the skull with dental cement. The cortical surface was kept moist with warm artificial cerebrospinal fluid (ACSF, 37 °C) throughout the experiment. ACSF solution was prepared with NaCl 120 mM, KCl 3.2 mM, K2HPO4 1 mM, HEPES 10 mM, NaHCO3 26 mM in a final volume of 1L of deionized water. pH was adjusted at 7.4 and the stock was stored at 4°. At the time of experiment, the solution was completed with CaCl2 2 mM and MgCl2 1 mM. LFP recordings were obtained using a silver/silver-chloride (Ag/AgCl) wire inserted into a borosilicate glass micropipette (tip diameter ∼1.5 µm, filled with ACSF). The electrode was advanced through a small opening in the coverslip to a depth of 250–300 µm (cortical layers 2/3) at a 45° angle using a motorised micromanipulator, positioned within the two-photon imaging field of view. Signals were amplified 1000× (EXT−02F amplifiers; NPI Electronics), band-pass filtered (0.1–1000 Hz), digitised at 10 kHz (16-bit), and line noise (50 Hz) removed with a notch filter.

Following acquisition of a baseline LFP recording of at least 3 min, 4-aminopyridine (15 µM dissolved in ACSF) and VU0463271 (10 µM in ACSF and 0.1% DMSO) were applied directly to the brain surface. Simultaneous two-photon imaging (830 nm excitation, 1 frame every 5 s) and LFP recordings were acquired continuously throughout drug application as a single time series. Imaging frames were temporally aligned to LFP recordings via shared acquisition timestamps generated by the end of frame signal. Seizures were identified following the criteria already described^5^; onset and offset were determined for each ictal event and fluorescence epochs were extracted accordingly. Mean ROI fluorescence traces were fitted with a logistic sigmoid function (seizure onset) and a decaying exponential function (seizure offset) using nonlinear least squares optimisation, from which the half-rise time and decay time constant (*τ*) were extracted.

## Statistical analysis

Statistical analyses were performed in MatLab. Data are presented as mean ± SEM or median ± MAD as indicated. The Wilcoxon signed-rank test was used for paired comparisons; the Mann-Whitney test was used for unpaired comparisons. The Kolmogorov-Smirnov test was used to compare cumulative distributions. Goodness of fit was assessed by R^2^. A significance threshold of p<0.05 was used throughout unless otherwise stated.

## Code availability

All custom computer code and algorithms used to generate the results and analyze the imaging data reported in this study are available in a GitHub repository. The code is organized into sections corresponding to the different experimental analyses.

## Data availability

The primary numerical data underlying all graphical representations, statistical analyses, and plots generated during this study (Figures 1–5 and Supplementary Figures) will be made available as a Source Data file upon publication. The raw datasets generated and/or analyzed during the current study are available from the corresponding authors upon reasonable request due to their large file size.

## AI-assisted writing

AI-assisted support was provided by Claude (Sonnet 4.6, Anthropic) during manuscript preparation, including assistance with scientific writing, grammar, reference identification, and methodological review. All scientific content, data, interpretations, and conclusions are the sole responsibility of the authors.

## Supplementary Information

### Measurement of bleed through

The emission spectra of mClover3* and mCRISPRed partially overlap. Full separation of their fluorescence would require emission filters with very narrow bandwidths, which would be detrimental to signal intensity. However, bleed-through can be corrected by linear decomposition, as described in Sulis Sato et al. (2017^4^). The bleed-through-corrected fluorescence of mClover3* and mCRISPRed is given by:

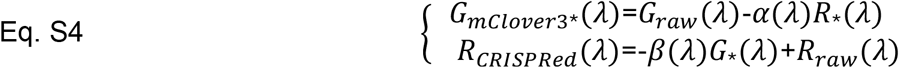

where *G*_*raw*_(*λ*) and *R*_*raw*_(*λ)* are the background-subtracted fluorescence signals measured in the green and red channels respectively, and *G*_*mClover3\**_(*λ)* and *R*_*CRISPRed*_(*λ)* are the corresponding bleed-through-corrected fluorescence values. In the main text, these are indicated as *g* and *r* for brevity. The bleed-through coefficient *α(λ*) represents the fractional contribution of mCRISPRed emission to the green channel, and *β(λ)* represents the fractional contribution of mClover3* to the red channel. To determine *α(λ)* and *β(λ)* for both one- and two-photon excitation, MEF cells were transfected separately with mClover3* or mCRISPRed and imaged using the same set-up as subsequent experiments.

Fluorescence was measured in both channels across a range of excitation wavelengths and the coefficients were computed as described in Supp. Fig. 2. It should be noted that these coefficients depend not only on the intrinsic spectral properties of the sensor but also on the specific imaging set-up, as they are influenced by detector gain, detector spectral sensitivity, and the spectral transmittance of the emission filters. To monitor the longitudinal stability of these parameters, the relative gain of the two channels should be verified by repeated measurement of a stable fluorescence standard such as a diluted solution of a fluorophore^4^. Since bleed through depends on the relative gains and detection efficiency of the two channels, it is different for each set up employed in this study.

**Supplementary Figure 2.**
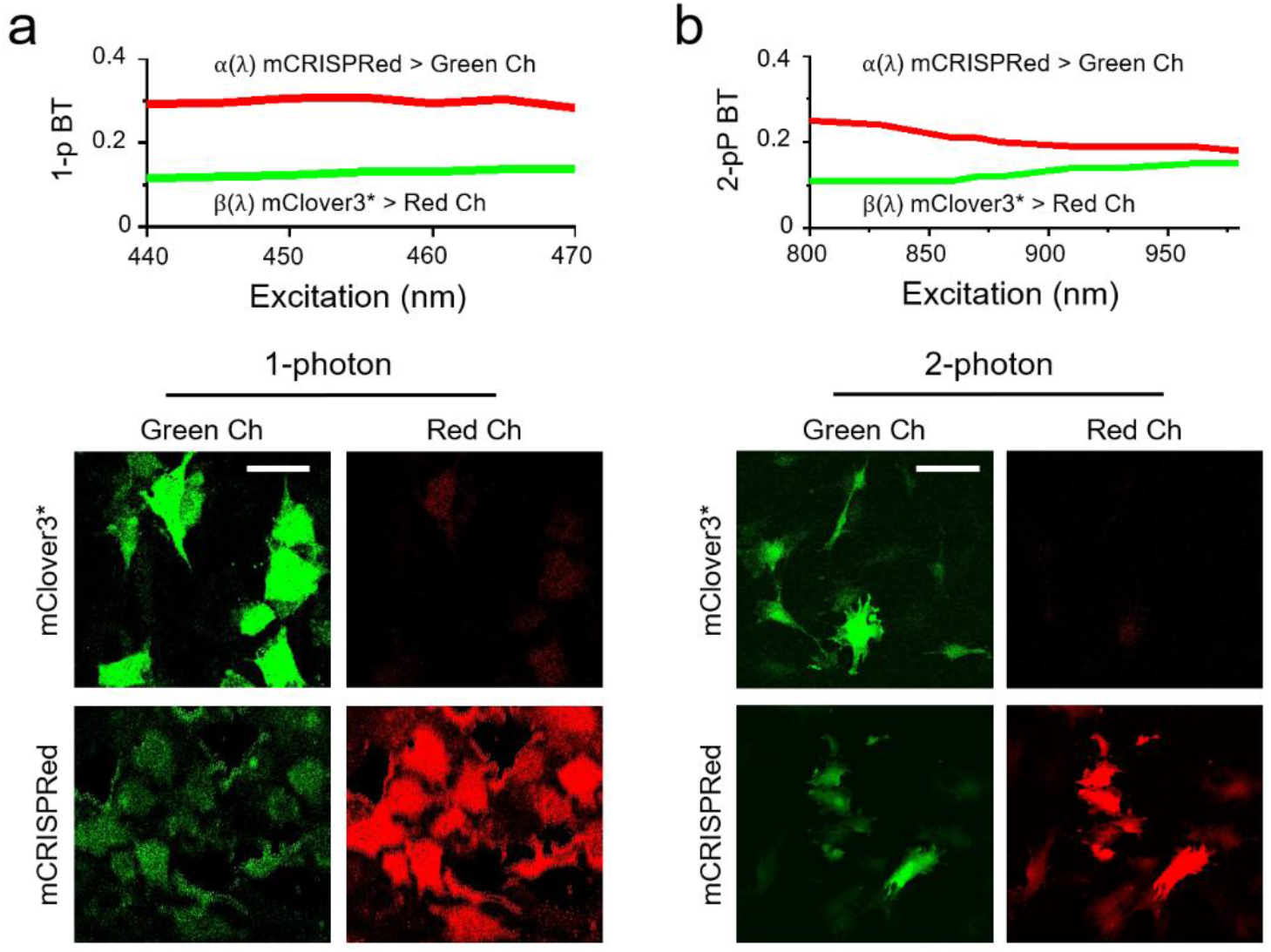
Measurement of bleed-through coefficients. **a**) Representative images MEF cells expressing mClover3* or mCRISPRed individually were imaged using one-photon excitation on a Leica Stellaris confocal microscope equipped with a white-light laser. Scale bar: 40 µm. Similar measurements were performed across all imaging set-ups used in the three participating laboratories. **b**) Equivalent measurements performed using two-photon excitation. Scale bar: 50 µm. To avoid saturation artefacts, only cells whose fluorescence signal remained within the bottom 75% of the detector dynamic range were included in the analysis.

**Derivation of the equation 1**

Equation 1 describes the dependence of θ on pH. It differs from the heuristic equation used in Sulis Sato et al. 2017^4^ in that it is derived from an explicit model of the biochemical equilibrium between three states of iClima in the presence of Cl^−^ and H^+^, illustrated in Fig. 1a and described here:

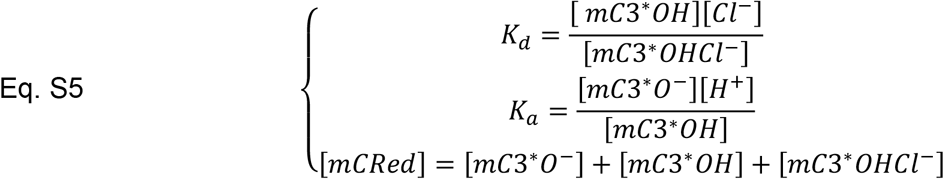

where *mC3^∗^O^−^, mC3^∗^OH* and *mC3^∗^OHCl^−^* denote the deprotonated, protonated, and protonated-chloride-bound states of mClover3 respectively, mCRed denotes mCRISPRed, and *K*_*a*_ and *K*_*d*_ are the acid dissociation constant and the chloride dissociation constant respectively. The conservation equation is based on the 1:1 stoichiometric relationship between mClover3* and mCRISPRed imposed by the fusion design. Solving the system for the mClover3* species gives:

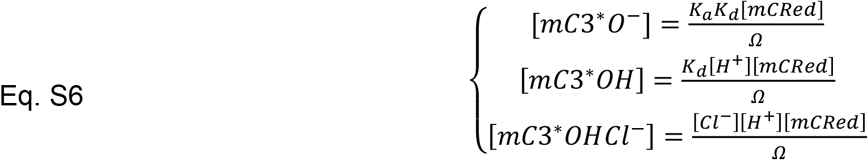

where we define:

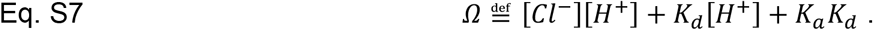

The model rests on the following assumptions:

1. The dependence of the excitation spectrum of mClover3* on pH arises from the protonation state of a single residue, as established for the green moiety of LSSmClopHensor^35^. These states are denoted *mC3^∗^OH and mC3^∗^O^−^* respectively
2. Cl^−^ binding to mC3*O*^*−*^ is forbidden (*i.e*. the dissociation constant for this reaction is infinite), as for the green moiety of LSSmClopHensor^36^, so that Cl^−^ can only bind to *mC3^∗^OH* to form the complex *mC3^∗^OHCl^−^*. The complex *mC3^∗^OHCl^−^* is assumed to be non-fluorescent.
3. *mC3^∗^OH and mC3^∗^O^−^* have the same emission spectrum. Only small changes occur in the pH range 5.0–9.0 (emission peak shifting from 510 to 523 nm) due to excited-state proton transfer (ESPT) from the chromophore phenol to a proton-acceptor residue^36^.
4. mCRISPRed excitation and emission spectra are independent on Cl^-^ and pH.

The fluorescence intensity collected from a fluorophore *X* under two-photon excitation at wavelength *λ* is given by:

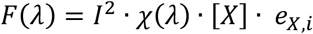

where *I* is the excitation intensity, *χ* is the two-photon excitation cross-section spectrum, [X] is the fluorophore concentration, and *e*_*i*_ is a collection factor accounting for emission spectrum, spatial integration over the point-spread function, emission filter transmittance, detector spectral response, and photon collection efficiency. The signal collected in the green channel after dark subtraction and bleed-through correction is therefore:

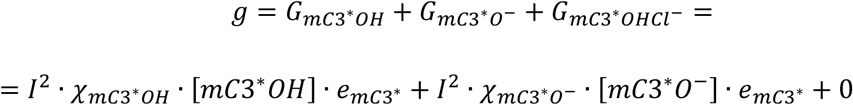

which, using assumption 2 (*mC3^∗^OHCl^−^* is non-fluorescent) and substituting from Eq. S6, simplifies to:

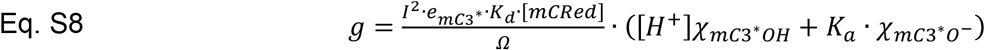

where e_mC3_∗ is the collection factor of mClover3* in the green channel. Analogously:

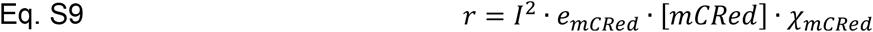

where r is the signal measured in the red channel after dark subtraction and bleed-through correction and *e*_*mCRed*_ is the collection factor of mCRISPRed in the red channel. The g/r ratio is therefore:

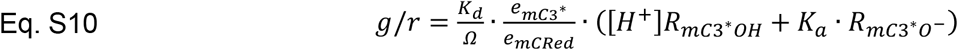

Where *R*_*mC3*_∗_*OH*_ and *R*_*mC3*_∗_*O*_− are the excitation spectra of the protonated and deprotonated forms of mClover3^*^ normalized by the excitation spectrum of mCRISPRed. Note that *g/r* is the quantity shown in Figure 1d. Equation S10 can be rewritten as:

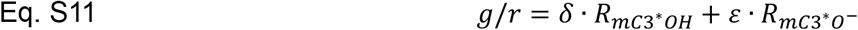

Thus, the *g/r* ratio at any pH and [Cl^−^] is a linear combination of the spectra of the protonated and deprotonated states of mClover3*. Representing δ and ε as coordinates on a Cartesian plane (Figure 1e), we define the angle:

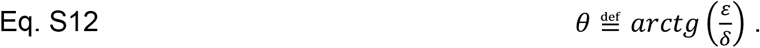

Substituting *δ* and ε, the equation simplifies to:

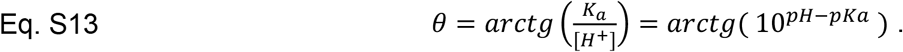

Note that, while δ and ε depend on both pH and [Cl^-^], *θ* only depends on pH and pK_a_. This is the same approach used in Sulis Sato et al 2017^4^, but here we are using the fully protonated and fully deprotonated spectra as a basis for the linear decomposition. The advantage of this approach is that it provides a theoretical function with just one parameter (pK_a_) to fit the experimental data (*i.e., θ* measured at different pH) instead of a heuristic sigmoidal function with many parameters. In practice, measuring excitation spectra at pH values extreme enough to approximate full protonation and full deprotonation is not experimentally feasible, as the protein would denature. We therefore use spectra measured at two accessible pH values — pHa (“acidic”) and pHb (“basic”) — as basis functions to decompose *g/r* :

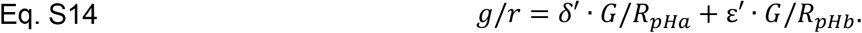

Here *δ′* and ε′ have an apex since they are different in nature by the coefficients appearing in Eq S11. Note again that, differently from the quantity *R*, which is the ratio of the excitation spectra and it is hard to measure experimentally, here the bases *g/r*_*pHa*_ and *g/r*_*pHb*_ are actually the ratio between green and red fluorescence measured experimentally and shown in figure 1d. However, it does not prevent this theoretical approach, since we can express *g*_pH1_ and *g*_pH2_ as a function of the protonated and deprotonated spectra:

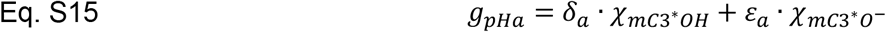

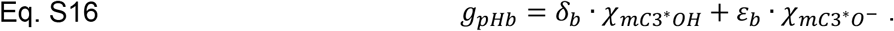

Substituting Eqs. S15 and S16 into Eq. S14:

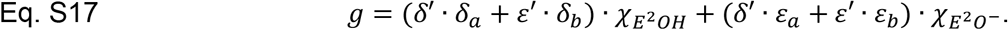

Defining 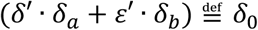 and 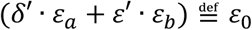 Eq. S17 Is seen to be equivalent to Eq S11. From the definitions of ε and δ it follows that:

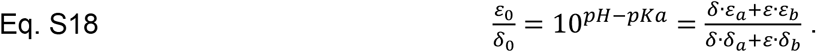

This last equation can be rearranged as follows:

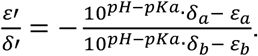

Isolating *δ*_*a*_ at the numerator and *δ*_*b*_ at the denominator:

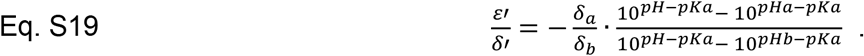

We now focus on the factor 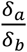. From the definition 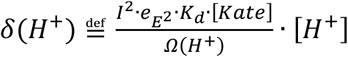

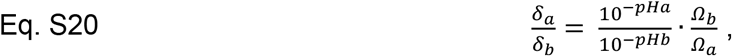

with *Ω*_*a*_ = [*Cl*^−^]10^−*pHa*^ + *K*_*d*_10^−*pHa*^ + *K*_*a*_*K*_*d*_ and *Ω*_*b*_ = [*Cl*^−^]10^−*pHb*^ + *K*_*d*_10^−*pHb*^ + *K*_*a*_*K*_*d*_. Since Ω also depends on [Cl^-^]i, it is convenient to measure *g/r* at pHa and pHb in a medium with zero Cl^-^. With [*Cl*^−^] = 0:

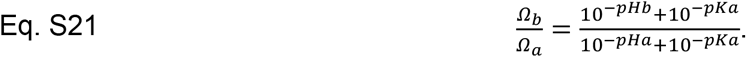

Substituting Eq. S21 into Eq. S20 and then into Eq. S19, we obtain:

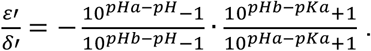

Which finally gives: Eq. S22

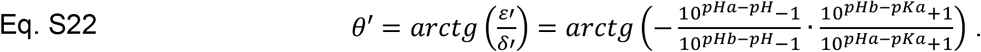

Eq S22 was used in two ways:

- During calibration.
  1. Spectra were measured at pH_a_=6 and pH_b_=8.
  2. The *g/r* ratio was measured at a series of known pH values (Fig. 1d) and fitted with Eq. S14, yielding a set of (*δ*^′^, *ε*^′^) pairs and thus a *θ* value for each pH.
  3. The *θ* values were fitted with Eq. S22 to obtain the pK_a_.

- To measure pH.
  1. The *g/r* ratio was measured in a sample at unknown pH.
  2. The ratio was fitted with Eq. S14 to obtain *δ*^′^, *ε*^′^ and thus *θ*.
  3. Eq S22 was inverted to yeld pH:

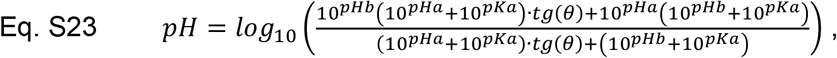

### Dynamics two-photon imaging of [Cl^-^]i during epileptic activity *in vivo*

We assessed the sensor functionality for extended imaging sessions designed to capture Cl^−^ fluxes during epileptic activity. Mice expressing iClima in the occipital cortex were anaesthetised with urethane. Focal epileptic seizures were induced by local application of 4-aminopyridine (4-AP) and VU0463271, following a protocol similar to Pracucci et al.^5^. Neural activity was monitored through simultaneous local field potential (LFP) recordings and two-photon imaging at the isosbestic wavelength (830 nm), enabling direct correlation of Cl^−^ optical signals with electrophysiological activity (Supp Fig. 3a–e).

Seizure episodes, clearly identifiable in the LFP recordings, were accompanied by a marked decrease in green fluorescence, consistent with a rise in [Cl^−^]i. The signal recovered upon seizure termination (Supp Fig. 3c–e). Analysis of individual seizure events revealed that [Cl^−^]i reached values of approximately 40 mM. Seizures of this magnitude cause intracellular acidification of approximately 0.2 pH units^4,12,37^, which we do not correct for in our Cl^−^ estimate. As shown in Fig. 1h, a pH shift of this magnitude introduces an error in the Cl^−^ estimate of approximately 7%, confirming that iClima provides reliable measurements even without simultaneous pH correction.

To quantify the kinetics of Cl^−^ transients, the g/r fluorescence ratio was extracted for each imaged neuron around seizure onset and offset separately (Supp Fig. 3f). To obtain metrics describing the onset of the Cl^−^ transient and its recovery, individual onset traces were fitted with a logistic function and individual offset traces with a single exponential decay; the same fitting approach is used in the analysis of Figures 4 and 5. Averaged across n=15 neurons from 1 mouse — encompassing 77 seizure onset episodes and 42 offset episodes, the lower number of the latter reflecting instances in which recovery was interrupted by a subsequent ictal event — the fits yielded a half-rise time of 2.6 s and a decay time constant of τ=18 s. The relatively slow post-ictal recovery likely reflects not only the kinetics of KCC2-mediated Cl^−^ extrusion but also the impaired transporter driving force during the post-ictal period, when extracellular K^+^ accumulation reduces the K^+^ gradient that drives cotransport^38,39^.

Taken together, these experiments demonstrate that iClima reliably reports [Cl^−^]i dynamics *in vivo*, resolving both the amplitude and kinetics of Cl^−^ transients during pathophysiological activity. The combination of high Cl^−^ affinity, pH robustness, and photostability makes iClima uniquely suited to capturing the complex ionic landscape of seizure activity, opening new avenues for investigating the role of Cl^−^ dysregulation in epilepsy and related disorders.

**Supplementary Figure 3.**
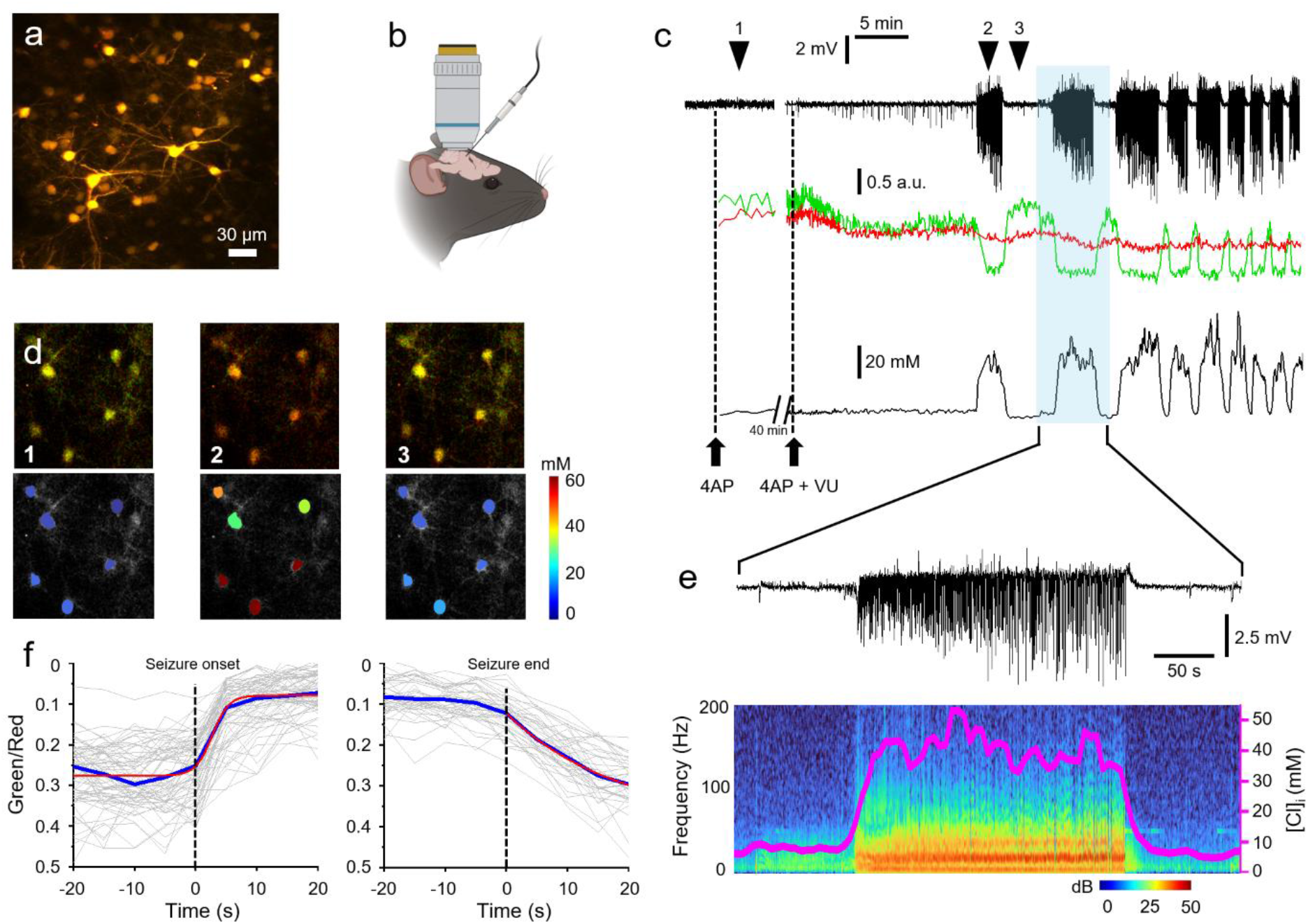
Dynamics of an epileptic seizure studied with iClima. **a**) Representative two-photon field of view acquired in vivo showing pyramidal neurons expressing iClima. **b**) Schematic of the simultaneous two-photon imaging (830 nm, isosbestic wavelength) and local field potential (LFP) recordings. 4-aminopyridine and VU0463271 were superfused onto the cortex at the indicated times. Generated with Biorender.com **c**) Top: LFP recording showing several ictal events. Middle: green and red fluorescence signals mediated on all neurons in the field after correction for dark offset and bleed-through. Bottom: estimated [Cl^−^]i, showing rapid increases during each ictal event followed by gradual and complete recovery. **d**) Top row: frames acquired at the time points indicated in panel c (left, baseline; middle, ictal event; right, post-seizure). Bottom: colour-coded estimates of [Cl^−^]i values (scale bar indicated). **e**) Expanded view of the highlighted ictal event, illustrating the temporal relationship between electrophysiological activity (LFP raw trace, top; spectrogram, bottom) and average [Cl^−^]i changes across imaged neurons (magenta line, n=15 neurons). **f**) Expanded view of [Cl^−^]i dynamics in pyramidal neurons (n=15 neurons, 77 rising and 42 recovery episodes of critical activity seizures from 1 mouse) around seizure onset and offset. Thin lines indicate individual neuronal time courses; the blue line is the median and red lines are the logistic and exponential fits used to extract kinetic parameters.

### Interference of the screen luminance on the imaging of Cl^-^ transients

The Cl^-^ transients represent a small modulation of the detected fluorescence and a potential concern in these experiments is that photons from the visual stimulation monitor could enter the microscope optical path and contaminate the detected fluorescence signal. To correct for this, movies were acquired under identical stimulation conditions with the excitation laser blocked, and the resulting dark signal was subtracted from the imaging data. To verify that this correction was sufficient, we repeated the visual stimulation protocol with a mechanical shutter inserted in the laser path upstream of the microscope entrance. The shutter was controlled by an Arduino board and alternated between open and closed states on a frame-by-frame basis, so that every second frame was acquired with the laser completely blocked (Supp. Fig. 4a). This scheme allowed the dark signal to be measured simultaneously with the real acquisition, on a per-ROI and per-frame basis, rather than relying on separate dark recordings.

Visual stimulation consisted of an equiluminant gray screen followed by five repetitions of a 5 s drifting grating (95% contrast) interleaved with 3 s gray periods, followed by a post-stimulus recovery epoch. ROIs were drawn around pyramidal cell somata and the dark-frame signal was subtracted on a frame-by-frame basis for each ROI (Supp. Fig. 4b). Robust visually evoked responses persisted after this correction (Supp. Fig. 4c), confirming that the observed fluorescence changes reflect genuine biological signals rather than stray light contamination.

To quantify the maximum possible contribution of light contamination to the ratiometric signal, we constructed artificial time series for each ROI and channel as the sum of the measured baseline fluorescence F (assumed constant, i.e. no chloride-dependent signal) and the dark time series D(t). These artificial traces represent the signal that would be expected if the only source of variation were fluctuations in the dark signal. Analysis of these artificial cells revealed ΔR/R_0_ values negligibly small compared to the real responses (Supp. Fig. 4d), with a temporal profile entirely different from that of genuine visually evoked responses (Supp. Fig. 4e).

**Supplementary Figure 4.**
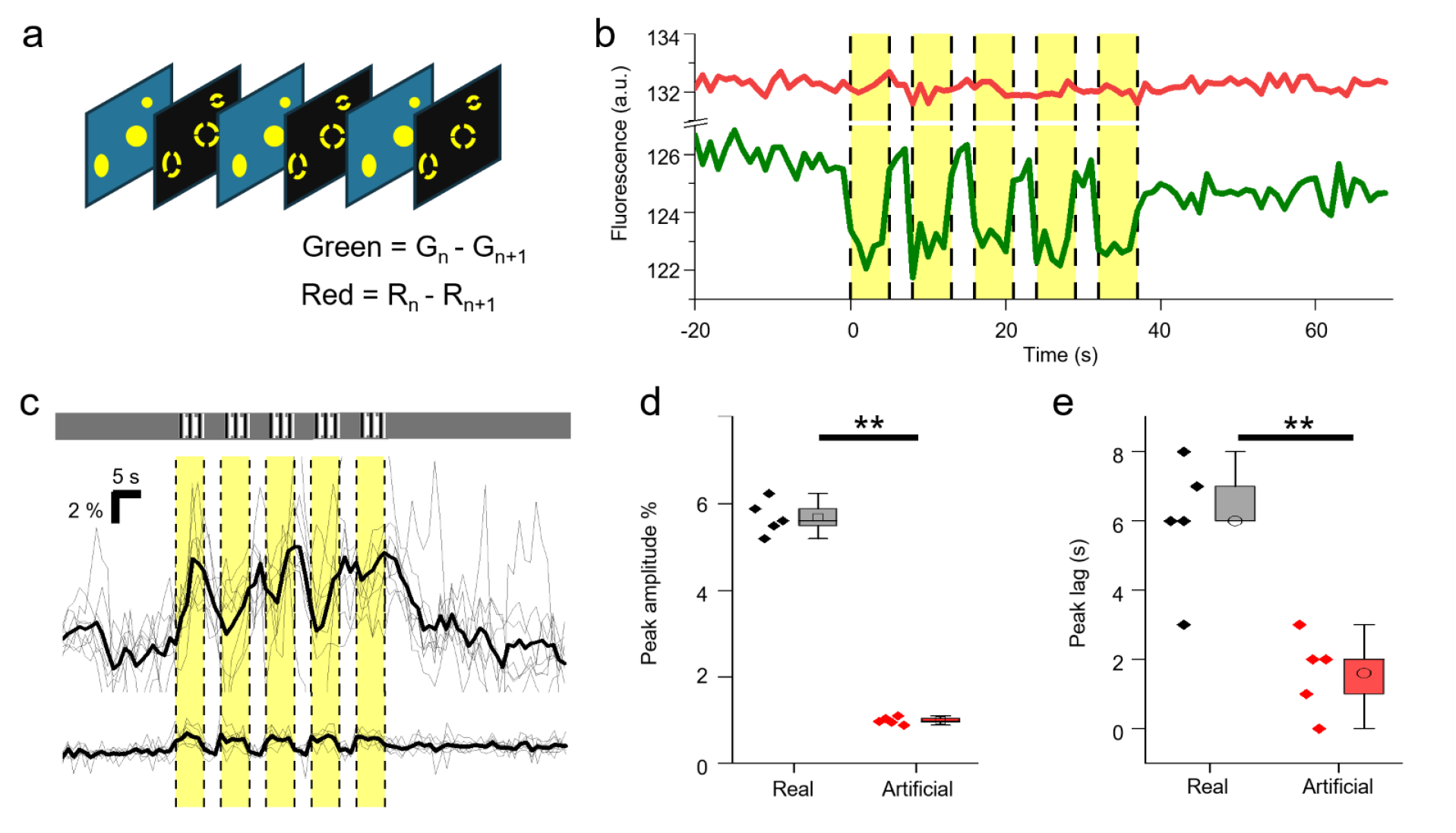
Control experiments confirming the specificity of visually evoked iClima signals. **a**) Schematic of the acquisition protocol used to assess and correct for potential light contamination from the visual stimulation monitor. A mechanical shutter was inserted in the excitation laser path and opened and closed alternately on a frame-by-frame basis, yielding interleaved “light” and “dark” frames. **b**) Representative dark-frame fluorescence traces (green and red channels) from a single ROI during visual stimulation (yellow shading). The dark-frame signal was subtracted on a per-ROI, per-frame basis. **c**) Population-averaged ΔR/R_0_ responses to five repetitions of a 5 s drifting grating after dark-frame subtraction. Top: real cells; bottom: artificial cells constructed from baseline fluorescence plus dark signal (see text). Thick line: mean; thin lines: individual trials. The persistence of robust responses in real cells after correction confirms that detected signals are not attributable to stray light from the monitor. **d**,**e**) Comparison of response peak amplitude (d) and temporal profile (e) between real cells and artificial cells, demonstrating the negligible contribution of light contamination to the ratiometric signal.

### Kinetics of the Cl transients *in vivo*

The Cl^−^ transient evoked by a brief sensory stimulus was fitted with a piecewise model. The rising phase was described by a logistic function, reflecting the progressive recruitment of GABAA receptors and the gradual increase in intracellular Cl^−^. At stimulus offset, the current through GABAA receptors terminates and recovery is dominated by KCC2-mediated Cl^−^ extrusion, which in the linear approximation follows a single exponential decay:

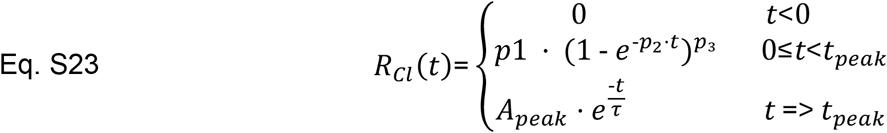

where *p*_1_ is the response amplitude, *p*_2_ is the rate constant of the rising phase, *p*_3_ is a shape parameter, *tpeak* is the time of peak response, *τ* is the time constant of the Cl^-^ recovery. *A*_*peak*_ is the fitted amplitude reached at *t*_*peak*_. The model was fitted to the data shown in Fig. 4f and provided an excellent description of the observed responses. Fit parameters for all four mice are reported in Table 1. To assess goodness of fit, we tested two hypotheses on the residuals. First, if the model captures the response accurately, residuals during the response phase should be statistically indistinguishable from residuals during the baseline period, which reflect noise alone; this was confirmed by a Mann-Whitney test. Second, the mean residual across the response period should not differ from zero; this was confirmed by a t-test. Both tests were passed for all four mice (Table 1).

**Table S1.**
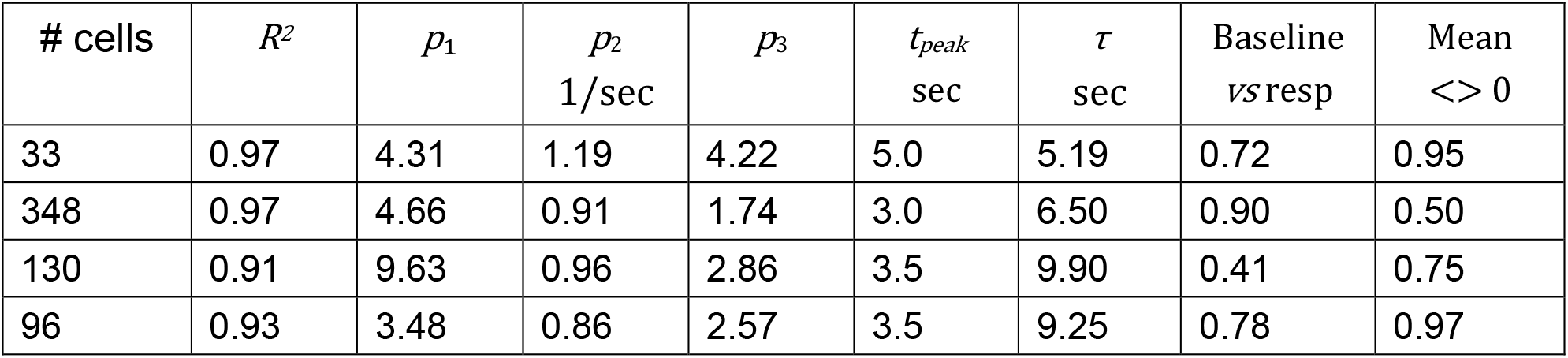
Summary of fit parameters for the average traces from four mice shown in Fig. 4f. The last two columns reports the p values of the statistical testing performed on the fit residuals.

### Combined Ca^2+^ and Cl^−^ imaging: linear separation of GCaMP6f and iClima signals

Separation of the Ca^2+^- and Cl^−^-dependent signals exploits the absence of a Ca^2+^-dependent response when iClima is excited at 830 nm (Supp. Fig. 5). While it is possible to subtract the 830 nm Cl^−^ transient directly from the 920 nm mixed signal, this approach would amplify noise by a factor of 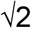 since the two channels are acquired independently. We therefore followed a model-based strategy, in which a noiseless analytical model of the Cl^−^ response is fitted to the 830 nm data and subtracted from the 920 nm signal on a cell-by-cell basis.

**Supplementary Figure 5.**
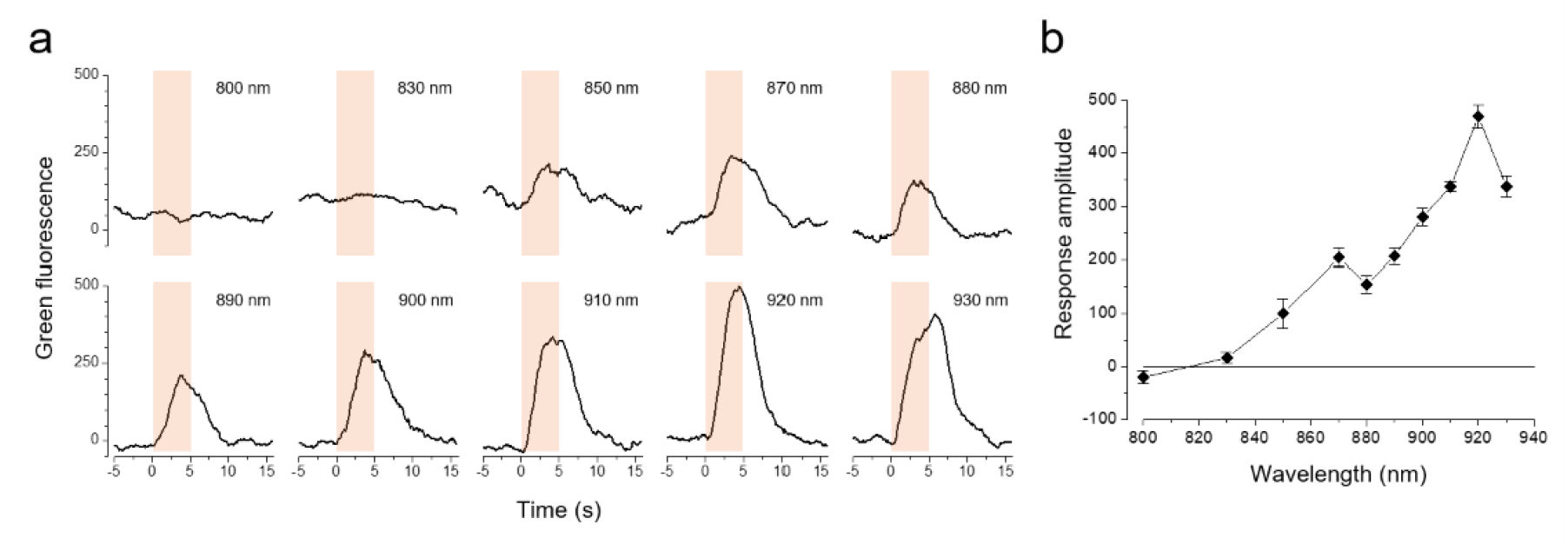
Isosbestic point of GCaMP6f. **a**) Dynamics of the green fluorescence in response to a 5 s long drifting grating at different excitation wavelengths. The shaded area shows the timing of the stimulus. **b**) Amplitude of the response as a function of the excitation wavelength.

For each imaged cell, the Cl^−^ transient recorded at 830 nm was described by a piecewise model *M*_830_(t), defined in Eq. S23. The model was fitted independently to the 830 nm response of each cell and approximates the data well (Supp. Fig. 6a). Since the *ΔR/Rbaseline* response of iClima to a Cl^−^ change arises from static quenching, the amplitude and kinetics of the Cl^−^ response should in principle be identical at 830 nm and 920 nm, provided that the Cl^−^-insensitive baseline signal is estimated correctly at both wavelengths. However, this condition cannot be met here because GCaMP6f contributes to the baseline fluorescence at both wavelengths, introducing an error in *R*_*baseline*_. Crucially, this error affects only the amplitude of the Cl^−^ response, not its kinetics. The 830 nm model was therefore scaled by a single free parameter γ and fitted to the 920 nm response:

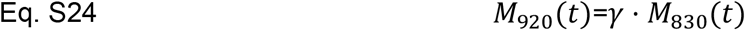

*γ* was estimated by minimising the sum of squared residuals between the 920 nm signal and the scaled 830 nm model. To prevent the faster Ca^2+^ transient from biasing the fit, only the late phase of the 920 nm response, after the Ca^2+^ transient has decayed, was used for the estimation of γ (Supp. Fig. 6b). After scaling and subtraction of the Cl^−^ component, the corrected green fluorescence signal at 920 nm was used to compute ΔF/F, yielding the isolated Ca^2+^ response shown in Fig. 5.

**Supplementary Figure 6.**
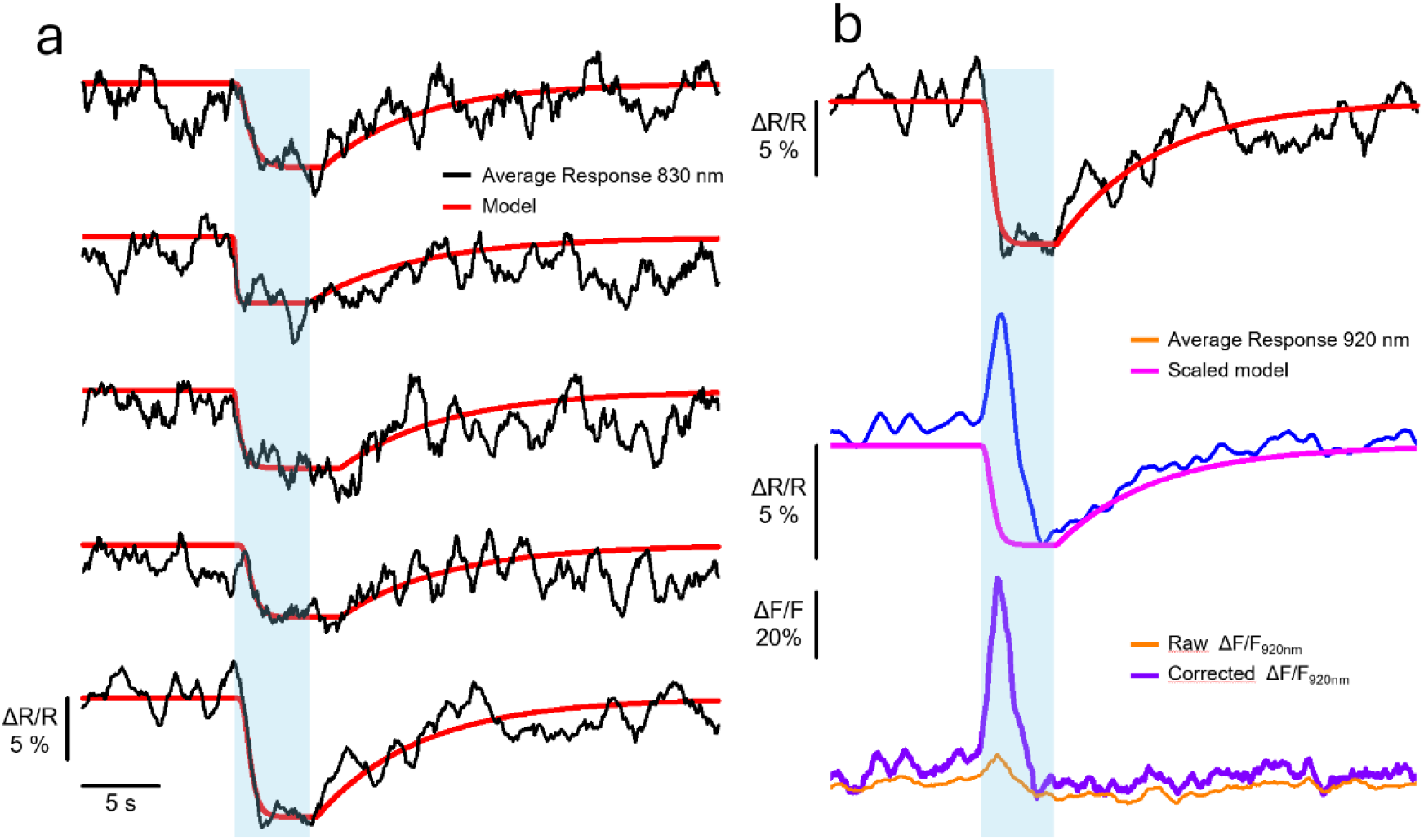
Linear separation of Ca^2+^ and Cl^−^ signals. **a**) Cl^−^ transients expressed as the percentage change of the G/R ratio in response to a 5 s drifting grating. The response is directed downward since green fluorescence decreases as [Cl^−^]i increases. Representative traces from 5 neurons recorded in 2 mice are shown; the red line shows the piecewise model fit from Eq. 23. The shaded area indicates the stimulus presentation period. **b**) Responses recorded at 920 nm excitation contain both the Cl^−^ and Ca^2+^ transients. The scaled 830 nm model (Eq. S24) was fitted to the late phase of the 920 nm response to estimate the scaling factor γ, excluding the early phase to prevent the faster Ca^2+^ transient from biasing the fit. After scaling and subtraction of the Cl^−^ component, the corrected green fluorescence at 920 nm was used to compute ΔF/F, yielding the isolated Ca^2+^ response.

